# Polo-like kinase Cdc5 orchestrates Cdk1 regulation via Swe1 and Mih1 during meiotic prophase I exit

**DOI:** 10.64898/2025.12.23.696139

**Authors:** Sara González-Arranz, Isabel Acosta, Irene Gil-Torres, Ethel Queralt, Beatriz Santos, Pedro A. San-Segundo

## Abstract

Defects in chromosome synapsis and meiotic recombination activate a checkpoint that, in budding yeast, delays exit from meiotic prophase I by inhibiting the Ndt80-dependent expression of regulators including the cyclin *CLB1* and the polo-like kinase (PLK) *CDC5*. Additionally, Swe1-mediated inhibitory phosphorylation of cyclin-dependent kinase 1 (Cdk1/Cdc28) reinforces this arrest. Once the checkpoint is released, Cdk1 activation is essential for meiosis I entry and requires removal of inhibitory phosphorylation on tyrosine 19, governed by the opposing activities of the Swe1 kinase and the Mih1 phosphatase. Here, we dissect how this network is rewired at the prophase I to meiosis I transition. We show that Swe1 is essential for checkpoint maintenance, but not for its initial activation. We also demonstrate that Cdc5 promotes Cdk1 activation through a dual mechanism: by inducing Swe1 degradation and facilitating Mih1 nuclear translocation. Unlike in mitosis, Cdc5-dependent Swe1 degradation in meiosis does not require CDK-mediated priming and can occur when both proteins are artificially colocalized, indicating a distinct regulatory mode. Our findings uncover a novel function for Cdc5 in promoting meiotic cell cycle progression beyond its known roles in recombination and synaptonemal complex disassembly, highlighting how conserved cell cycle regulators are adapted to meiosis.

## INTRODUCTION

Meiosis is a specialized cell division that produces haploid gametes from diploid precursor cells, ensuring genetic diversity and genome stability across generations. This process consists of one round of DNA replication followed by two consecutive rounds of chromosome segregation, meiosis I and meiosis II. A defining feature of meiosis is the programmed induction of DNA double-strand breaks (DSBs) during prophase I by the conserved topoisomerase-like enzyme Spo11 ^1^. These DSBs initiate homologous recombination (HR), a repair process that facilitates the formation of crossovers (COs) between homologous chromosomes, which are essential for proper chromosome alignment and segregation ^2^. Meiotic recombination is closely coordinated with chromosome synapsis, the intimate association between paired homologous chromosomes mediated by the synaptonemal complex (SC). The SC is a highly ordered proteinaceous structure composed of two lateral elements (LEs) and a central region. In *Saccharomyces cerevisiae*, the LEs are primarily formed by Hop1, Red1, and Rec8-containing cohesin, while the transverse filament protein Zip1 constitutes the major structural component of the central region, along with the central element components Ecm11 and Gmc2 ^3^. Synapsis is required for the proper maturation of interhomolog COs, which, in turn, ensure accurate chromosome segregation.

To preserve genomic integrity, cells have evolved surveillance mechanisms known as checkpoints. In meiotic cells, the meiotic recombination checkpoint (MRC) prevents progression to meiosis I until recombination is successfully completed. The MRC is activated in response to persistent DSBs or synapsis defects, leading to the activation of a signaling cascade that delays the transition from prophase I to meiosis I. The MRC consists of several functional modules: sensors, mediators, effectors, and targets. Checkpoint activation occurs when unrepaired DSBs accumulate, as observed in *S. cerevisiae* mutants defective in synapsis (i.e., *zip1Δ*) or recombination (i.e., *dmc1Δ*) ^4,5^. The *zip1Δ* mutant lacks a functional SC, resulting in impaired synapsis and incomplete recombination that activate the checkpoint ^6-8^. On the other hand, recombinase-deficient *dmc1Δ* cells fail to promote interhomolog strand invasion, leading to an accumulation of unrepaired DSBs and checkpoint activation ^9^.

DNA resection at DSB ends generates single-stranded DNA (ssDNA) regions, which are rapidly coated by Replication Protein A (RPA). When DSB repair fails, this RPA-ssDNA complex recruits the Mec1 kinase (ATR in mammals) through its regulatory subunit Ddc2 (ATRIP in mammals) ^10,11^. The checkpoint signal is further amplified by Rad24 and the 9-1-1 complex (Ddc1-Rad17-Mec3), which promote Mec1 activation and downstream signaling ^12,13^.

A key event in checkpoint activation is the Mec1-dependent phosphorylation of Hop1 at T318, which facilitates the recruitment and activation of Mek1, the central effector kinase of the MRC, in a process that also involves Red1 ^14-17^. The AAA+ ATPase Pch2 plays a crucial role in modulating Hop1 conformation and is required to achieve proper levels of Hop1-T318 phosphorylation under checkpoint-inducing conditions ^6^. Active Mek1 phosphorylates and inhibits the Ndt80 transcription factor, preventing the expression of genes necessary for meiotic progression, including *CDC5* and *CLB1* ^18-21^ (Fig. 1a).

**Fig 1.**
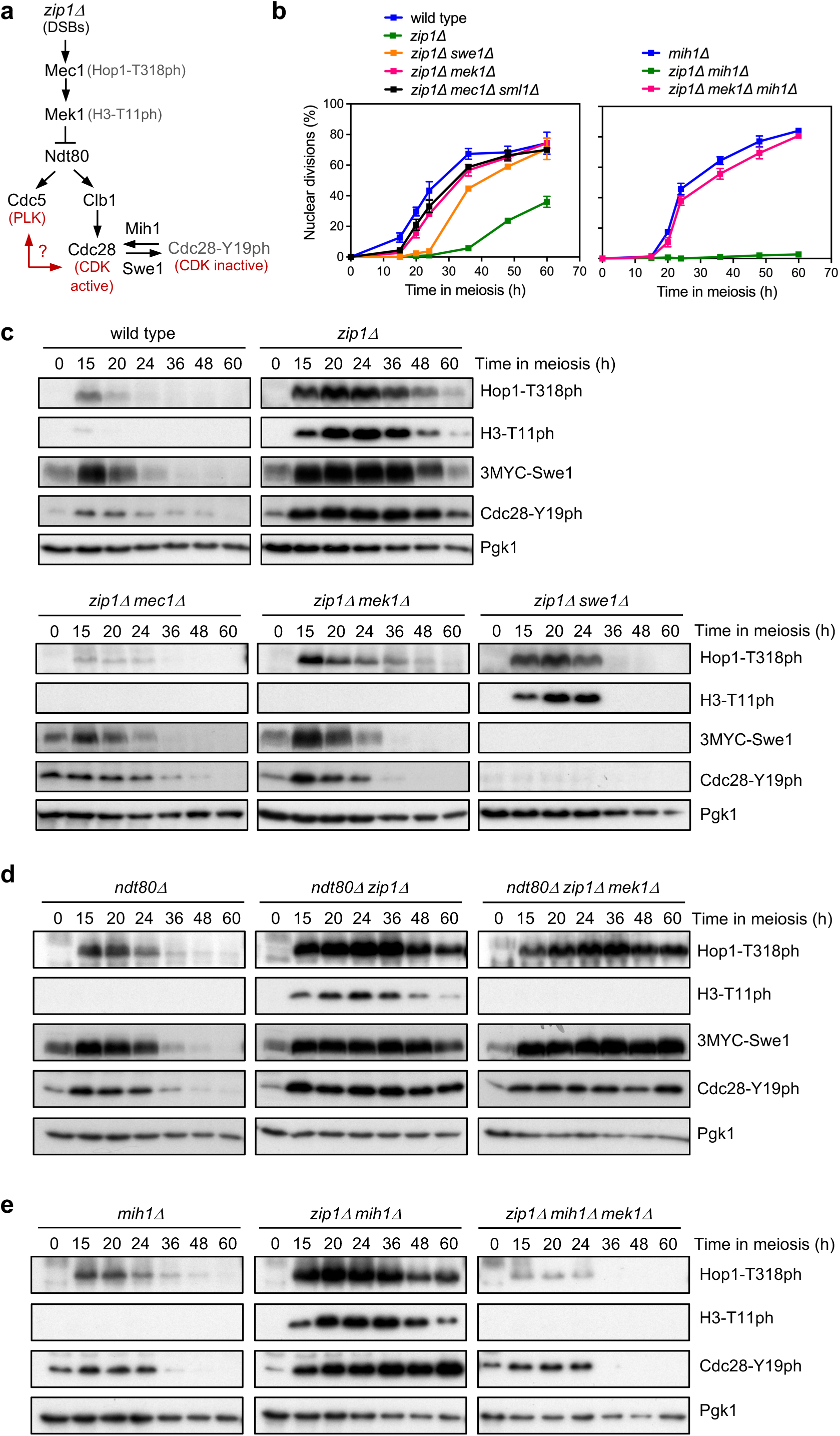
Regulation of Cdc28-Y19 phosphorylation by Swe1 and Mih1 during the meiotic recombination checkpoint response in *zip1Δ*. **(a)** Simplified schematic representation of the meiotic recombination checkpoint pathway in *S. cerevisiae* highlighting the proteins most relevant to this study. Phosphorylation reporters for Mec1, Mek1 and Swe1 are indicated in grey. The question mark denotes the potential crosstalk between PLK and CDK regulation explored here. **(b)** Time-course analysis of meiotic nuclear divisions; the percentage of cells containing two or more nuclei is represented. Error bars: SD; n=3. At least 300 cells were scored for each strain at every time point. (**c-e)** Western blot analysis of Mec1 and Mek1 checkpoint signaling (Hop1-T318ph and H3-T11ph, respectively), Swe1 production (detected with anti-MYC antibody), and Cdc28-Y19ph levels during meiotic time courses of the indicated strains. Pgk1 was used as loading control. Strains in (b-e) are DP1353 (wild type), DP1354 (*zip1Δ*), DP1407 (*zip1Δ mec1Δ sml1Δ*), DP1393 (*zip1Δ mek1Δ*), DP1157 (*zip1Δ swe1Δ*), DP1402 (*mih1Δ*), DP1403 (*zip1Δ mih1Δ*), DP1404 (*zip1Δ mek1Δ mih1Δ*), DP1398 (*ndt80Δ*), DP1397 (*ndt80Δ zip1Δ*) and DP1396 (*ndt80Δ zip1Δ mek1Δ*).

Cdc5 is the sole member of the Polo-like kinase (PLK) family in *S. cerevisiae* and serves as a crucial regulator of meiotic progression, promoting key events such as SC disassembly and the resolution of recombination intermediates ^22-25^. Its expression is tightly controlled by the transcription factor Ndt80, linking MRC deactivation to meiotic progression. Among several Ndt80 targets, Cdc5 is particularly important for driving the transition from prophase I to meiosis I ^20^. Like in mitosis, the order and timing of meiotic events is orchestrated by cyclin-dependent kinase (CDK) activity formed by a catalytic subunit (Cdc28 in *S. cerevisiae*, also known as Cdk1) and different cyclins ^26,27^. Clb1 is the major B-type cyclin required for meiosis I ^28,29^; therefore, checkpoint-dependent inhibition of Ndt80 ensures that both M-CDK and (PLK) activities remain low, maintaining meiotic arrest until recombination is successfully completed.

Beyond cyclin binding, Cdk1 activity is also regulated by inhibitory phosphorylation. In *S. cerevisiae*, the Swe1 kinase (homologous to WEE1) phosphorylates Cdc28 at Tyr19 (equivalent to Tyr15 in *Schizosaccharomyces pombe* Cdc2 and mammalian CDK1), reducing its activity. This inhibitory phosphorylation is counteracted by the Mih1 phosphatase (homologous to CDC25), which promotes Cdk1 activation at the G2/M transition ^30-32^. Unlike in fission yeast and mammals, Cdc28-Tyr19 phosphorylation (Cdc28-Y19ph) does not play a major role in unperturbed mitotic progression in budding yeast ^30,33^. However, it is critical for restraining mitotic entry in response to the morphogenesis checkpoint, which delays the cell cycle when budding fails ^34^. In meiosis, the primary target of the MRC is the Ndt80 transcription factor, but Swe1-dependent inhibition of Cdc28 additionally contributes to the cell cycle delay imposed by the checkpoint ^35,36^. Genetic evidence suggests that these two regulatory mechanisms: (1) Mek1-dependent inhibition of Ndt80 and, consequently, Clb1 production, and (2) Swe1-dependent inhibition of Cdc28, function through distinct branches of the checkpoint response pathway, although they are likely functionally interconnected ^37^ (Fig. 1a).

While Swe1’s role in the MRC has been long recognized, the molecular mechanisms governing its regulation during meiosis, as well as those of its counteracting phosphatase Mih1, remain unclear. To address this, we investigated the regulation of inhibitory Cdc28-Y19ph in response to checkpoint activation triggered by the absence of Zip1. Here, we reveal that Swe1 is required for the maintenance, but not the initial activation, of the MRC in *zip1Δ* cells. Furthermore, we found that, although Mih1 is dispensable for meiotic progression under unperturbed conditions, its absence strengthens the checkpoint-dependent meiotic arrest of *zip1Δ* cells. Using an inducible *CDC5* allele, we demonstrate that Cdc5 kinase activity drives Swe1 degradation even when both proteins are artificially colocalized at an ectopic cellular location. Additionally, we show that Cdc5 not only regulates Cdc28-Y19 phosphorylation through Swe1 but also promotes Mih1 nuclear entry at the prophase I to metaphase I transition, facilitating Cdc28-Y19 dephosphorylation. Overall, our findings uncover a previously unrecognized role of the Cdc5 polo-like kinase in coordinating the downregulation of inhibitory CDK phosphorylation by targeting both positive and negative regulators of M-CDK activity. Thus, our results highlight how multiple regulatory layers integrate to ensure timely meiotic progression.

## RESULTS

### Swe1 is required to maintain, but not to establish, checkpoint activity

The involvement of Swe1 in the MRC response has been previously established, but the precise molecular mechanisms that regulate its activity during meiosis, along with those of its antagonist, the phosphatase Mih1, are still poorly understood. To address this, we investigated the regulation of Cdc28 inhibitory phosphorylation at Tyr19 in response to checkpoint activation triggered by the absence of the Zip1 protein.

To monitor Swe1 levels under different meiotic conditions, we generated strains expressing epitope-tagged Swe1. Although a C-terminally 3HA-tagged Swe1 version has been used in other mitotic studies, we found that this version did not support meiotic recombination checkpoint function (Fig. S1). However, an N-terminally tagged Swe1 version with three copies of the MYC epitope was fully functional (Fig. S1), and we used this construct for subsequent analyses. To characterize the checkpoint response in *zip1Δ* cells, we examined Swe1 levels alongside its target, Cdc28-Y19 phosphorylation (ph), and two established checkpoint activation markers: Hop1-T318 phosphorylation by Mec1, and histone H3-T11 phosphorylation by Mek1. Meiotic division kinetics were assessed by DAPI staining of nuclei.

The *zip1Δ* mutant exhibited a pronounced meiotic delay, coinciding with high levels of Hop1-T318ph and H3-T11ph. These checkpoint markers decreased only at later time points, correlating with the resumption of meiotic progression in a fraction of *zip1Δ* cells. Swe1 and Cdc28-Y19ph levels also followed a similar temporal pattern (Fig. 1b, c).

As expected, deletion of *MEC1* (in a *sml1Δ* background) or *MEK1* almost completely suppressed the *zip1Δ*-induced delay, restoring nearly wild-type kinetics of meiotic nuclear divisions and wild-type dynamics of Swe1 protein levels (Fig. 1b,c). Consistent with the sequential action of both checkpoint kinases, Hop1-T318ph was negligible in *zip1Δ mec1Δ* cells, while H3-T11ph was abolished in both *zip1Δ mec1Δ* and *zip1Δ mek1Δ* cells. In contrast, the *zip1Δ swe1Δ* double mutant retained strong phosphorylation of Hop1-T318 and H3-T11 and exhibited meiotic arrest at early time points. However, after 24 hours in meiosis, Hop1-T318ph and H3-T11ph sharply declined, and nuclear divisions resumed (Fig. 1b,c).

These findings indicate that Swe1 is not required for the initial establishment of Mec1-Mek1 checkpoint signaling in response to meiotic defects caused by *zip1Δ*, but it is essential for sustaining checkpoint activity.

### Phosphorylation of Swe1 at S385 (SQ site) is dispensable for the meiotic recombination checkpoint

Although earlier reports excluded Swe1 from participating in the DNA damage response during the mitotic cell cycle ^33,38^, more recent studies have revealed that Swe1 functions in a parallel branch to Rad53 in the checkpoint response to replicative stress. Furthermore, phosphorylation of Swe1 at its unique S/T-Q consensus site for the Mec1 kinase (Ser385) is required for proper S-phase checkpoint function ^39^. Since Swe1 appears to act downstream of Mec1 in the meiotic recombination checkpoint (MRC) (see above), we investigated whether phosphorylation of Swe1 at its Ser385 Mec1 target site is also necessary for the MRC. To do this, we mutated Ser385 to alanine to generate the *swe1-S385A* phosphomutant. In contrast to the *zip1Δ swe1Δ* mutant (Fig. 1b, Fig. S1), the *zip1Δ swe1-S385A* double mutant failed to sporulate and exhibited a significant delay in meiotic progression, similar to the *zip1Δ* single mutant (Fig. S2a, S2b). The meiotic block in *zip1Δ swe1-S385A* was dependent on the MRC, as it was relieved by deletion of *MEK1*, leading to the production of predominantly inviable spores (Fig. S2a, S2b). Thus, we conclude that, unlike the replication checkpoint in mitotic cells, phosphorylation of Swe1 at its unique SQ Mec1 consensus site is not involved in the MRC response.

### High levels of Swe1 persist in *zip1Δ ndt80Δ*-arrested cells independently of Mek1

To assess the impact of meiotic cell-cycle progression on Swe1-dependent Cdc28-Y19 phosphorylation, we performed western blot analysis of the checkpoint markers described above in an *ndt80Δ* mutant background, which prevents exit from prophase I. Curiously, despite being arrested in prophase I, the *ndt80Δ* single mutant exhibited a decline in Swe1 accumulation (and consequently, Cdc28-Y19ph) after the 24-hour time point (Fig. 1d). This observation suggests that cell-cycle progression is not required for Swe1 elimination under otherwise unperturbed conditions. In contrast, checkpoint activation in *zip1Δ ndt80Δ* cells, as indicated by increased phosphorylation of Hop1-T318 and H3-T11, resulted in sustained levels of Swe1 and Cdc28-Y19ph throughout the entire time course (Fig. 1d). Notably, unlike in *zip1Δ mek1Δ* (Fig. 1c), both Swe1 and Cdc28-Y19ph remained elevated even in the absence of upstream checkpoint signaling in *ndt80Δ zip1Δ mek1Δ* cells (Fig. 1d). These findings suggest that, at least under checkpoint-inducing synapsis-defective conditions caused by the absence of Zip1, an Ndt80-dependent event is required for Swe1 removal and the subsequent relief of its inhibitory effect on CDK.

### Cdc5 is required to remove Swe1-dependent Cdc28-Y19 phosphorylation under checkpoint conditions

We previously developed a conditional system that allows sequential activation and deactivation of the MRC by controlling *ZIP1* expression from the *GAL1* promoter with β-estradiol ^40^. This setup enables precise monitoring of exit from checkpoint-induced prophase I arrest and entry into meiosis I (Fig. 2a). Using this inducible *ZIP1-GFP-IN* system, we observed that the Ndt80-dependent induction of Cdc5 production upon checkpoint deactivation revealed by the drop in H3-T11 phosphorylation, closely correlates with a sharp decrease in Swe1 levels and the rapid loss of Cdc28-Y19 inhibitory phosphorylation (Fig. 2b; Fig. S3a). This observation suggests that Cdc5 may downregulate Swe1 function. In mitotic cells, several lines of evidence indicate that recovery from the arrest imposed by the morphogenesis checkpoint involves Cdc5-dependent degradation of Swe1 at the bud neck ^41-44^. Given that *CDC5* expression is a pivotal event launched by Ndt80 to promote entry into meiosis I, we investigated whether Cdc5 is responsible for counteracting the inhibitory effect of Swe1 on CDK activity established under checkpoint conditions to drive meiotic progression as checkpoint signaling diminishes.

**Fig 2.**
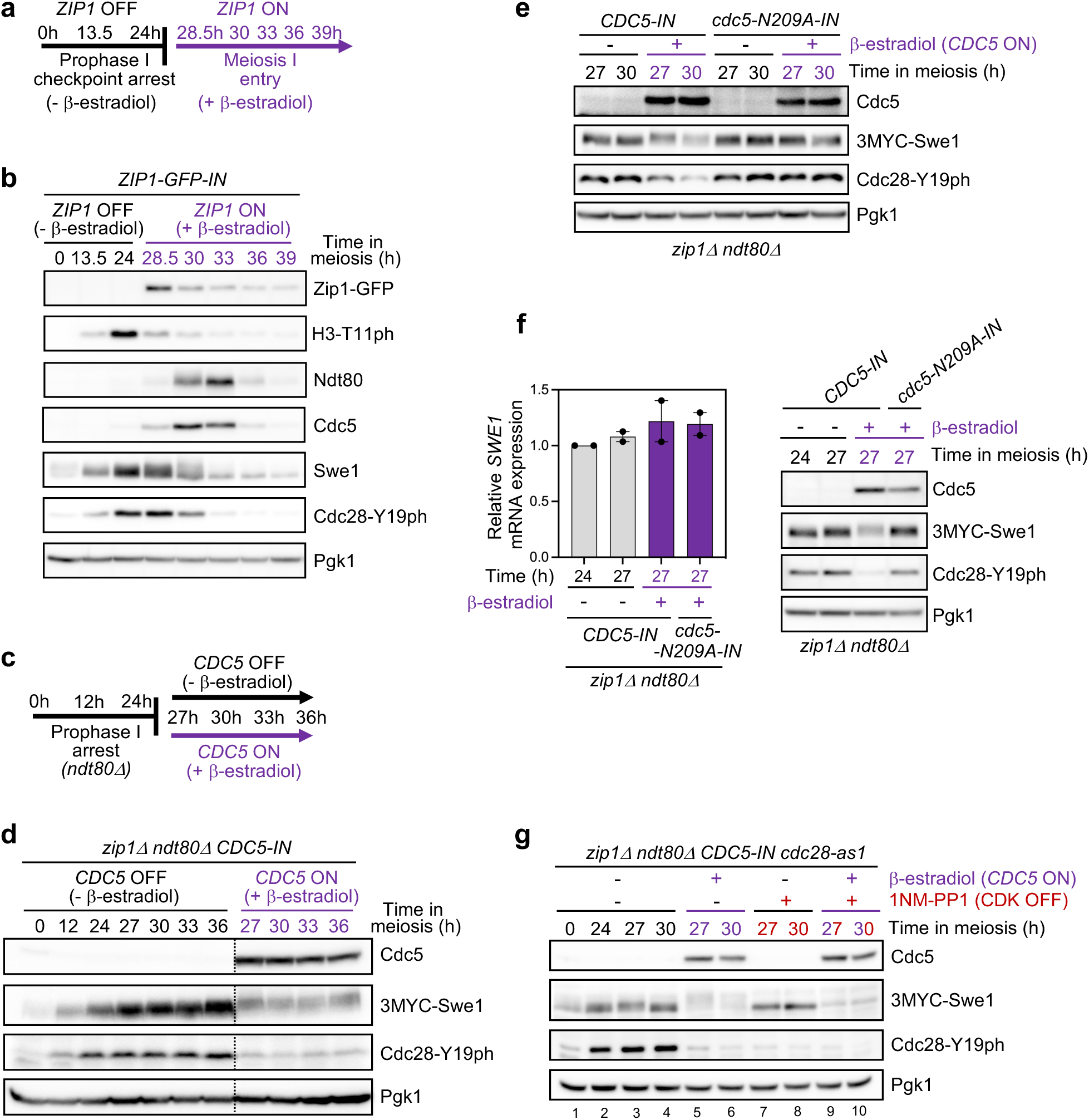
Cdc5 kinase activity promotes Swe1 degradation independent of CDK. **(a)** Experimental setup used to sequentially activate and inactivate the checkpoint in an inducible *ZIP1-GFP-IN* strain. In the absence of β-estradiol (*ZIP1* OFF) the lack of *ZIP1* triggers the checkpoint. Addition of β-estradiol (1 μM) after 24 h in checkpoint-arrested cells induces *ZIP1-GFP* expression and the ensuing checkpoint inactivation as Zip1 is loaded onto chromosomes ^40^. **(b)** Western blot analysis of the indicated proteins during checkpoint deactivation as outlined in (a). The inducible *ZIP1-GFP-IN* strain is DP1185 (*P_GAL_-ZIP1-GFP GAL4-ER*). **(c)** Schematic representation of *CDC5* induction experiments in *CDC5-IN* strains. β-estradiol (1 µM), or ethanol as control, were added at 24 h in meiosis, when most *ndt80Δ* cells are arrested in prophase I, and samples were collected at different time points. **(d-e)** Western blot analysis of Cdc5 (or Cdc5-N209A) and Swe1 production, and Cdc28-Y19 phosphorylation. Pgk1 was used as loading control. **(f)**. The left panel shows absolute expression levels of *SWE1* mRNA, normalized to the invariant control gene *NUP85*, in *CDC5-IN* and *cdc5-N209A-IN* prophase I cells in the presence or absence of β-estradiol, as indicated. Expression was quantified by digital PCR (n=2). Individual values are plotted along with the mean and range. Quality control parameters of the digital PCR analysis are provided in Figure S4. The right panel shows western blot analysis of Cdc5, Swe1, and Cdc28-Y19ph protein levels from the same samples used for mRNA expression measurement. Strains are DP1711 (*zip1Δ ndt80Δ CDC5-IN*) and DP2121 (*zip1Δ ndt80Δ cdc5-N209A-IN*). **(g)** Western blot analysis of Cdc5 and Swe1 production, and Cdc28-Y19 phosphorylation in the presence/absence of *CDC5* expression and/or CDK activity, as depicted. Pgk1 was used as loading control. β-estradiol (5 µM), or ethanol as control, were added at 24 h in meiosis. The 1NM-PP1 analog (1 µM), or DMSO as control, were also added at 24 h in meiosis. Strains are DP1711 (*zip1Δ ndt80Δ CDC5-IN*), DP2121 (*zip1Δ ndt80Δ cdc5-N209A-IN*), and DP1932 (*zip1Δ ndt80Δ CDC5-IN cdc28-as1*). Quantification of relative protein levels for (b-g) is shown in Fig. S3.

To test this hypothesis, we utilized the *CDC5-IN* allele, which allows conditional expression of *CDC5* upon β-estradiol addition ^25^. We introduced *CDC5-IN* into a *zip1Δ ndt80Δ* strain, in which the absence of *ZIP1* triggers checkpoint activation, and *NDT80* deletion provokes permanent prophase I arrest. The *zip1Δ ndt80Δ CDC5-IN* strain was induced to enter meiosis and, at the 24-hour time point, when most cells in the culture are blocked in prophase I in the BR strain background, the culture was split in two. One half was supplemented with β-estradiol, while the other was treated with the solvent ethanol, and samples were collected at successive time points (Fig. 2c). Consistent with previous results (Fig. 1d), the *zip1Δ ndt80Δ CDC5-IN* mutant, in the absence of β-estradiol, maintained fairly constant levels of Swe1 and Cdc28-Y19 phosphorylation throughout the experiment (Fig. 2d; Fig. S3b). Remarkably, activation of *CDC5* expression led to a rapid and dramatic decrease in Swe1 protein levels. This reduction in Swe1 was mirrored by a decline in Cdc28-Y19ph. Interestingly, the residual Swe1 protein detected in the presence of Cdc5 exhibited slower gel mobility (Fig. 2d). Since this experiment was performed in an *ndt80Δ* background, we conclude that Cdc5 is the sole relevant target of Ndt80 responsible for downregulating Swe1 protein amount and reducing Cdc28-Y19ph under *zip1Δ* checkpoint conditions. To determine whether the kinase activity of Cdc5 is required to downregulate Swe1 and Cdc28-Y19ph, we generated an inducible version of *CDC5* carrying a mutation that abolishes its kinase activity (*cdc5-N209A-IN*). Induction of the kinase-dead Cdc5-N209A version had little or no effect on Swe1 protein and Cdc28-Y19ph levels (Fig. 2e; Fig. S3c). Moreover, measurement of *SWE1* mRNA levels in *CDC5-IN* and *cdc5-N209A-IN* strains revealed that they are not diminished upon *CDC5* induction (Fig. 2f; Fig. S3d, S4), ruling out the possibility that the reduction in Swe1 protein triggered by Cdc5 stems from lower *SWE1* gene expression. Thus, we conclude that a Cdc5-dependent phosphorylation event is responsible for Swe1 degradation.

### Reduction of Swe1 levels promoted by Cdc5 is independent of CDK activity

Polo-like kinase (PLK) action is often primed by prior CDK phosphorylation of substrates, creating Polo-box binding sites that recruit the kinase for subsequent phosphorylation ^45^. Moreover, Cdc28 activates Cdc5 in vegetative cells ^46,47^. To determine whether the downregulation of Swe1 by Cdc5 requires prior CDK priming, we utilized a *zip1Δ ndt80Δ CDC5-IN cdc28-as1* strain, allowing CDK inhibition by the ATP analog 1NM-PP1 ^48^, in the presence or absence of estradiol to control *CDC5* induction (Fig, 2G, Fig. S3e). In the absence of *CDC5* induction, Swe1 accumulated regardless of CDK activity status (lanes 1–4 and 7–8). However, upon CDK inhibition, Swe1 gel mobility collapsed to a more discrete band, accompanied by a drastic reduction in Cdc28-Y19 phosphorylation, despite the continued presence of Swe1 protein when Cdc5 is absent (lanes 7–8), indicating that Swe1 is inactive in this condition. Notably, *CDC5* induction led to a marked decrease in Swe1 levels and a concomitant reduction in Cdc28-Y19 phosphorylation, both in the presence (lanes 5–6) and absence (lanes 9–10) of CDK activity. Taken together, these results indicate (1) that the retarded gel migration of Swe1 upon *CDC5* induction largely depends on CDK activity, (2) that CDK is also involved in a feedback loop to promote the inhibitory action of Swe1, namely phosphorylation of Cdc28-Y19, and (3) that the effect of Cdc5 in driving Swe1 degradation is exerted independently of CDK activity, implying that CDK priming is not required.

### Cdc5 promotes nuclear clearance of Swe1 during meiotic prophase I exit

To analyze the subcellular localization of Swe1, we generated an N-terminal GFP-tagged version of the protein. To track the dynamics of GFP-Swe1 localization during meiosis, we performed time-lapse live-cell microscopy. Given the poor synchrony of our BR strain background, we adapted the *P_CUP1_-IME1*-based meiotic synchronization system ^49^ to increase synchrony in our cultures (Fig. S5a; see Methods section). We observed that GFP-Swe1 expressed from its native promoter produced a very weak fluorescent signal in meiotic prophase I cells (Fig. S5b). Thus, to enhance detection, we constructed a *P_HOP1_-GFP-SWE1* version, in which expression was driven by the strong meiotic prophase-specific *HOP1* promoter. In this case, GFP-Swe1 exhibited a more conspicuous signal, localizing predominantly to the nucleus during meiotic prophase I (Fig. S5b). This GFP-Swe1 version is functional in the meiotic recombination checkpoint, as *zip1Δ swe1Δ* cells transformed with a plasmid carrying *P_HOP1_-GFP-SWE1* sustained Cdc28-Y19 phosphorylation (Fig. S5c). Moreover, integration of *P_HOP1_-GFP-SWE1* at the *SWE1* genomic locus augmented the meiotic progression delay of *zip1Δ*, consistent with high levels of Swe1 kinase activity (Fig. S5d, S5e). Time-lapse microscopy experiments revealed that the nuclear GFP-Swe1 signal disappeared as cells transitioned from prophase I to meiosis I, as monitored by spindle pole body (SPB) separation using Spc42-mCherry as a marker (Fig. 3a; Video S1). To determine whether the disappearance of GFP-Swe1 is specifically triggered by Cdc5 rather than being solely a consequence of meiotic cell-cycle progression, we analyzed *zip1Δ ndt80Δ CDC5-IN P_HOP1_-GFP-SWE1* cells, which remain blocked in prophase I. In these cells, in addition to the diffused nuclear signal, we frequently observed a prominent nuclear or cytoplasmic GFP-Swe1 focus, likely reflecting the high protein accumulation caused by checkpoint induction and the strong expression of *P_HOP1_-GFP-SWE1* in *zip1Δ ndt80Δ* prophase I-arrested cells. Importantly, upon β-estradiol addition to induce *CDC5* expression, we observed a progressive reduction in nuclear GFP-Swe1, even in the absence of meiosis I entry (Fig. 3b, 3c), indicating that Cdc5 is the unique Ndt80 target required for Swe1 degradation. We conclude that Cdc5 removes nuclear Swe1 during exit from prophase I.

**Fig 3.**
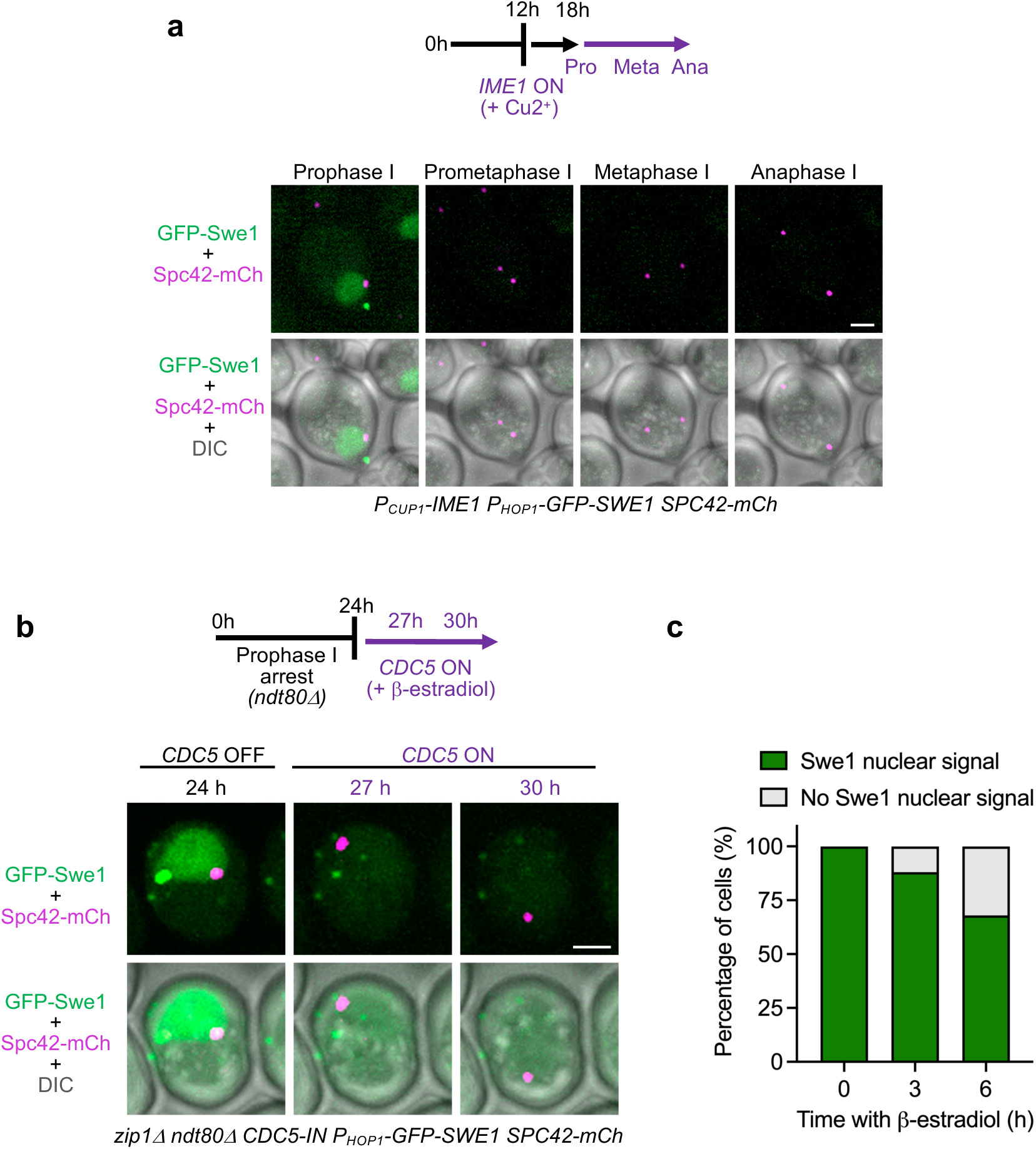
Cdc5 drives Swe1 nuclear removal at the prophase I to meiosis I transition. **(a)** Time-lapse fluorescence microscopy analysis of GFP-Swe1 dynamics during meiosis in *P_CUP1_-IME1*-synchronized cells. The spindle-pole body (SPB) marker Spc42-mCherry was used for staging. GFP-Swe1 goes away as SPBs separate. The experiment outline is presented on top. A representative individual cell is shown. Scale bar, 2 μm. The strain is DP1922 (*P_CUP1_-IME1 P_HOP1_-GFP-SWE1*) transformed with pSS449 (*SPC42-mCherry*). **(b)** Nuclear localization of GFP-Swe1 in *zip1Δ ndt80Δ* prophase I-arrested cells disappears as *CDC5* expression is induced by β-estradiol, as depicted on top. A representative cell with this behavior is shown. Scale bar, 2 μm. **(c)** Quantification of cells with GFP-Swe1 nuclear signal from the experiment shown in (b); 300 cells were scored from 3 independent captured fields. The strain is DP1934 (*zip1Δ ndt80Δ CDC5-IN P_HOP1_-GFP-SWE1*) transformed with pSS449 (*SPC42-mCherry*).

### Artificial tethering of Cdc5 and Swe1 reveals a direct role for Cdc5 in Swe1 degradation

To determine whether Cdc5 directly acts on Swe1, we artificially forced the colocalization of both proteins at an ectopic subcellular compartment outside the nucleus, specifically at the eisosomes on the plasma membrane (PM). Previously, we used the eisosomal protein Pil1 fused to the GFP-binding protein (GBP) to efficiently tether the GFP-tagged Pch2 meiotic checkpoint protein at the PM ^50^. Here, we tested whether GFP-Swe1 and GFP-Cdc5, both expressed during meiotic prophase I under the *HOP1* promoter, could similarly be recruited to the PM in cells expressing Pil1-GBP-mCherry. Fluorescence microscopy revealed that nuclear GFP-Swe1 was efficiently sequestered at the cell periphery, colocalizing with Pil1-GBP-mCherry (Fig. 4a). Likewise, while GFP-Cdc5 primarily accumulated at the nuclear envelope in control cells, it was also detected at the PM in the presence of Pil1-GBP-mCherry (Fig. 4b). However, Cdc5 tethering at the eisosomes was less efficient than that of Swe1, as a fraction of GFP-Cdc5 remained in nuclear aggregates. Nevertheless, a significant portion of GFP-Swe1 and GFP-Cdc5 was expected to be brought together at the PM when co-expressed in *PIL1-GBP-mCherry* cells (Fig. 4c). To assess whether this forced ectopic colocalization affects Swe1 stability, we analyzed Cdc5 and Swe1 protein levels by western blot in prophase I-arrested *zip1Δ ndt80Δ PIL1-GBP-mCherry P_HOP1_-GFP-SWE1* strains transformed with plasmids expressing GFP-tagged wild-type Cdc5 or the kinase-deficient Cdc5-N209A version, as well as GFP alone as control (Fig. 4d; Fig. S3f). We found that expression of *GFP-CDC5* (lanes 5–8) led to reduced levels of GFP-Swe1 compared to the control expressing either *GFP* alone (no Cdc5; lanes 1–4), or the kinase-dead mutant *GFP-cdc5-N209A* (lanes 9–12). These results indicate that Cdc5 can promote Swe1 degradation when both proteins are artificially forced into close proximity at an ectopic location, suggesting that the Cdc5 kinase directly targets Swe1.

**Fig 4.**
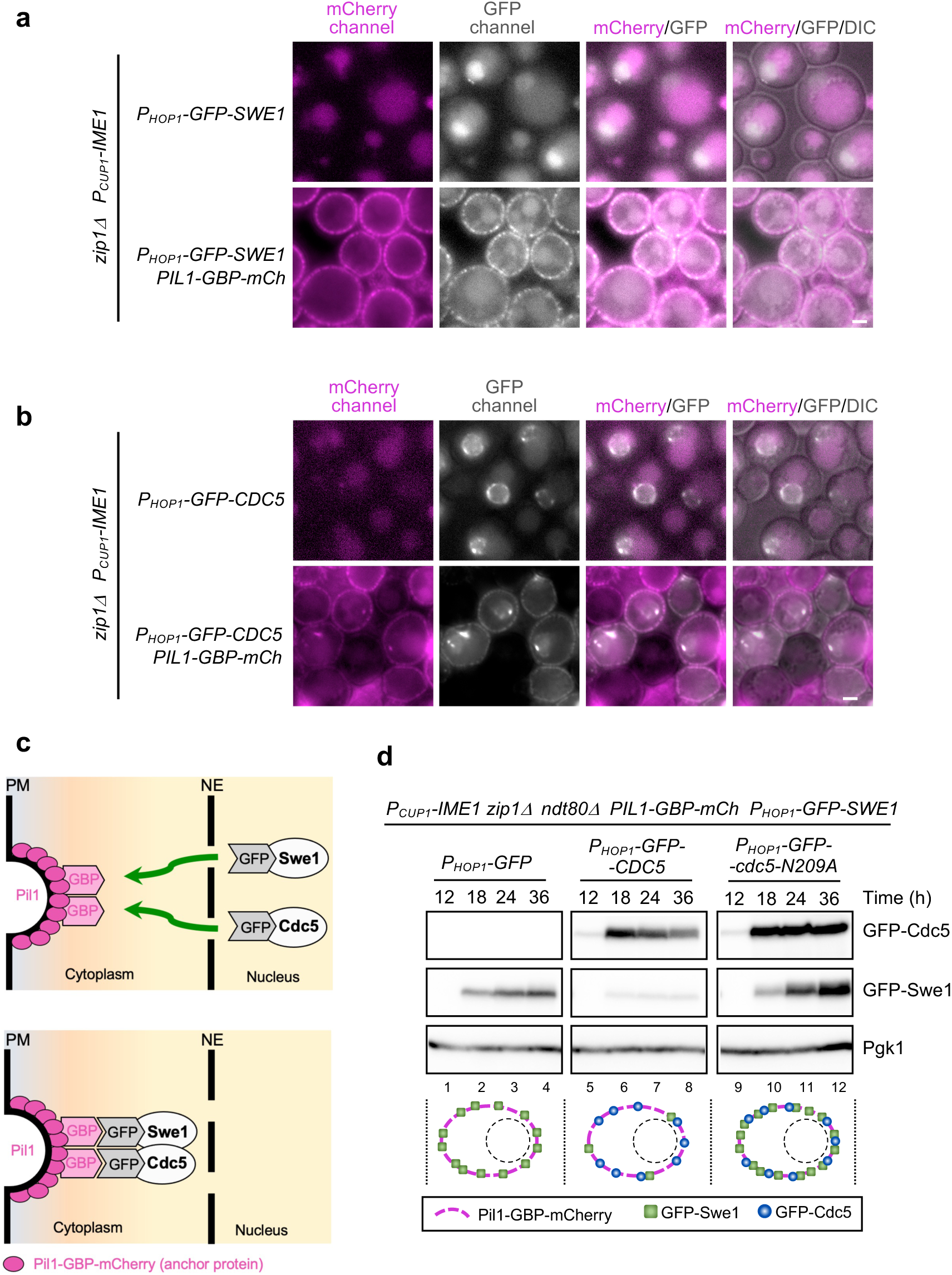
Forced ectopic colocalization of Cdc5 and Swe1 reduces Swe1 levels. **(a, b)** Fluorescence microscopy analysis of GFP-Swe1 (a), GFP-Cdc5 (b), and Pil1-GBP-mCherry distribution in meiotic cells of the indicated genotypes imaged 6 hours after Cu2^+^ addition in *P_CUP1_-IME1*-synchronized cultures. Fluorescence from mCherry and GFP is shown in magenta and grey, respectively. Top rows in (a) and (b) display control cells that do not express Pil1-GBP-mCherry. The diffuse magenta signal in these cells corresponds to background vacuolar autofluorescence. In contrast, Pil1-GBP-mCherry decorates the plasma membrane (bottom rows). Scale bar, 2 μm. Strains in (a) are DP1899 (*zip1Δ P_CUP1_-IME1 P_HOP1_-GFP-SWE1*) and DP1978 (*zip1Δ P_CUP1_-IME1 P_HOP1_-GFP-SWE1 PIL1-GBP-mCherry*). Strains in (b) are DP1935 (*zip1Δ P_CUP1_-IME1*) and DP2133 (*zip1Δ P_CUP1_-IME1 PIL1-GBP-mCherry*) both transformed with pSS441 (*P_HOP1_-GFP-CDC5*). **(c)** Cartoon depicting the artificial recruitment of GFP-Swe1 and GFP-Cdc5 to eisosomes containing Pil1-GBP. PM, plasma membrane; NE, nuclear envelope. **(d)** Western blot analysis of GFP-Cdc5 (detected with anti-Cdc5 antibody) and GFP-Swe1 (detected with anti-GFP antibody) production during *P_CUP1_-IME1*-induced meiotic time courses of the indicated strains. Pgk1 was used as loading control. The expected localization of Pil1-GBP-mCherry, GFP-Swe1, and GFP-Cdc5, or Cdc5, in each condition, according to panels (a) and (b) is illustrated below. Quantification of relative protein levels is shown in Fig. S3f. The strain used in (d) is DP2127 (*P_CUP1_-IME1 zip1Δ ndt80Δ PIL1-GBP-mCherry P_HOP1_-GFP-SWE1*) transformed with plasmids pSS248 (*P_HOP1_-GFP*), pSS441 (*P_HOP1_-GFP-CDC5*), or pSS484 (*P_HOP1_-GFP-cdc5-N209A*).

### The Mih1^CDC25^ phosphatase is required for resumption of meiotic progression in *zip1Δ*

To further investigate the regulation of CDK inhibitory phosphorylation in the *zip1Δ*-induced checkpoint, we explored the functional role of the Mih1^CDC25^ phosphatase, which counteracts Cdc28-Tyr19 phosphorylation by Swe1. The fact that deletion of *MIH1* in an otherwise wild-type background does not block meiosis and allows for the formation of viable spores led to the conclusion that Mih1 is not involved in the MRC response ^35^. Consistent with these findings, we observed that the *mih1Δ* single mutant displayed wild-type kinetics of meiotic progression and fairly normal dynamics of Cdc28-Tyr19ph, which largely disappeared after the 24-hour time point (Fig. 1b, 1e). However, deletion of *MIH1* in *zip1Δ* strains provoked a tighter meiotic block and the prolonged accumulation of high levels of Cdc28-Y19ph (Fig. 1b, 1e). This behavior contrasts with that of the *zip1Δ* single mutant, which, after a significant delay, eventually resumes meiotic divisions to some extent as Cdc28-Y19ph levels decrease at late time points (see above; Fig. 1b, 1c). Additionally, high levels of upstream checkpoint activation markers, Hop1-T318ph and H3-T11ph, were observed in *zip1Δ mih1Δ* cells (Fig. 1e), suggesting that the Mih1 phosphatase is required for checkpoint downregulation. Importantly, the robust meiotic arrest in *zip1Δ mih1Δ* was dependent on the MRC, as it was alleviated in *zip1Δ mih1Δ mek1Δ* cells (Fig. 1b), along with the persistence of Cdc28-Y19ph (Fig. 1e).

Thus, the Mih1 phosphatase is essential for the eventual attenuation of checkpoint activity in *zip1Δ* cells at later time points, allowing for the partial resumption of meiotic progression after a significant delay.

### Cdc5 triggers Mih1 nuclear entry during prophase I exit

To investigate the dynamics of Mih1 subcellular localization during meiosis, we performed time-lapse microscopy in *P_CUP1_-IME1*-synchronized cultures. We have previously observed that overexpression of *GFP-MIH1* in *zip1Δ* strains drastically reduces Cdc28-Y19 phosphorylation ^40^, indicating that N-terminal GFP tagging does not impair Mih1 phosphatase function. Since GFP-tagged Mih1 expressed from its native promoter was difficult to detect during meiosis, we integrated a *P_HOP1_-GFP-MIH1* construct at its endogenous locus to follow Mih1 localization in live cells. *P_HOP1_*-driven *GFP-MIH1* expression likely increases Mih1 levels, as *zip1Δ P_HOP1_-GFP-MIH1* cells progress in meiosis faster than *zip1Δ* with reduced Cdc28-Y19ph levels (Fig. S5d, S5d). Microscopy studies revealed that, during meiotic prophase I, Mih1 was predominantly cytoplasmic and largely excluded from the nucleus. As cells transitioned from prophase I to metaphase I, Mih1 progressively accumulated in the nucleus, coinciding with SPB separation. Then, the GFP-Mih1 signal gradually declined (Fig. 5a; Video S2).

**Fig 5.**
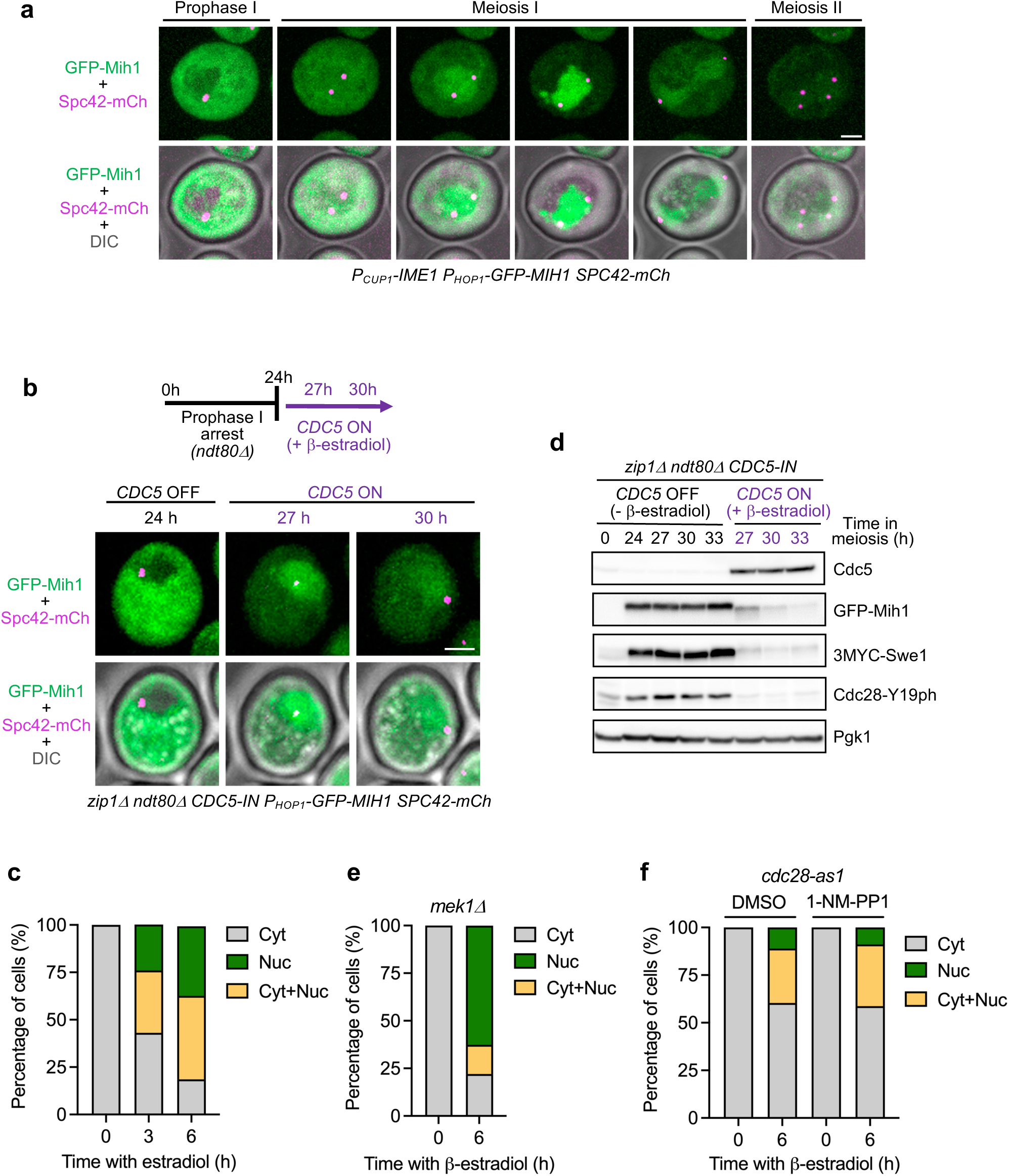
Cdc5 triggers nuclear entry of Mih1 at the prophase I to meiosis I transition. **(a)** Time-lapse fluorescence microscopy analysis of GFP-Mih1 dynamics during meiosis in *P_CUP1_-IME1*-synchronized cells, as outlined in Fig. 3a. The SPB marker Spc42-mCherry was used for staging. GFP-Mih1 translocates from the cytoplasm to the nucleus as SPBs separate. A representative individual cell is shown. Scale bar, 2 μm. The strain is DP1928 (*P_CUP1_-IME1 P_HOP1_-GFP-MIH1*) transformed with pSS449 (*SPC42-mCherry*). **(b)** Nuclear translocation of GFP-Mih1 in *zip1Δ ndt80Δ* prophase I-arrested cells as *CDC5* expression is induced by β-estradiol. A representative cell is shown. Scale bar, 2 μm. **(c)** Quantification of cells with GFP-Mih1 present in the cytoplasm (Cyt), nucleus (Nuc), and nucleus-plus-cytoplasm (Cyt+Nuc) from the experiment shown in (b). More than 600 cells were scored. **(d)** Western blot analysis of Cdc5, GFP-Mih1 and GFP-Swe1 production, and Cdc28-Y19 phosphorylation in the absence (*CDC5* OFF) or presence (*CDC5* ON) of β-estradiol. Pgk1 was used as loading control. Quantification of relative protein levels is shown in Figure S3G. The strain in (b-d) is DP1931 (*zip1Δ ndt80Δ CDC5-IN P_HOP1_-GFP-MIH1*) transformed with pSS449 (*SPC42-mCherry*). **(e)** Quantification of GFP-Mih1 subcellular distribution in *mek1Δ* cells analyzed as in (c). 300 cells were scored. The strain is DP1936 (*zip1Δ ndt80Δ CDC5-IN P_HOP1_-GFP-MIH1 mek1Δ*) transformed with pSS449 (*SPC42-mCherry*). **(f)** Quantification of GFP-Mih1 subcellular distribution in *cdc28-as1* cells, analyzed as in (c), treated with DMSO or with the 1NM-PP1 inhibitor, as indicated. 217 and 141 cells were scored for the DMSO and 1NM-PP1 conditions, respectively. The strain is DP1977 (*zip1Δ ndt80Δ CDC5-IN P_HOP1_-GFP-MIH1 cdc28-as1*) transformed with pSS449 (*SPC42-mCherry*).

To determine whether Cdc5 regulates Mih1 nuclear entry, we analyzed GFP-Mih1 localization in *zip1Δ ndt80Δ CDC5-IN P_HOP1_-GFP-MIH1 SPC42-mCherry* cells arrested in prophase I. In the absence of Cdc5 induction (no β-estradiol), GFP-Mih1 remained exclusively cytoplasmic in 100% of cells (Fig. 5b, 5c). Upon *CDC5* induction, Mih1 relocalized to the nucleus in ∼80% of cells, despite their persistent prophase I arrest manifested by a single Spc42-mCherry focus (Fig. 5b, 5c). At later time points, the GFP-Mih1 signal diminished (Fig. 5b). Western blot analysis revealed that Cdc5 expression induced faster-migrating Mih1 forms and a progressive reduction in Mih1 protein levels (Fig. 5d; Fig. S3g), suggesting that nuclear Mih1 is less stable.

It remained possible that Mih1 nuclear entry was an indirect consequence of checkpoint inactivation following Cdc5 induction. However, Mih1 relocalization was unaffected in the absence of the Mek1 checkpoint effector kinase (Fig. 5e), indicating that Cdc5 controls Mih1 independently of checkpoint signaling. We next assessed whether CDK activity is required for Mih1 relocalization by introducing the *cdc28-as1* allele. *CDC5* expression in *cdc28-as1* cells, in the absence of the ATP analog 1NM-PP1, promoted Mih1 nuclear entry in ∼40% of cells. This fraction remained unchanged when CDK activity was inhibited by 1NM-PP1 (Fig. 5f), indicating that Cdc5-mediated Mih1 redistribution does not require CDK activity. We note, however, that Mih1 nuclear entry is less efficient in *cdc28-as1* cells even without addition of the inhibitor.

Taken together, our results indicate that, in addition to promoting the nuclear disappearance of the Swe1 kinase, Cdc5 also facilitates Mih1 nuclear entry. These complementary actions contribute to the reduction of Cdc28-Y19 phosphorylation and the subsequent rise in CDK activity during the transition from prophase I to meiosis I.

## DISCUSSION

The successful transition from prophase I to meiosis I requires precise coordination between SC disassembly, the completion of meiotic recombination, and the initiation of chromosome segregation. This dependency is reinforced by the meiotic recombination checkpoint (MRC), which ensures that meiotic DSBs are fully repaired before progression by inhibiting key cell cycle regulators, including CDK and PLK. However, the precise mechanisms through which the MRC modulates these kinases to control meiotic progression remain incompletely understood. In this study, we elucidate how the polo-like kinase Cdc5 fine-tunes CDK activity by regulating Cdc28 phosphorylation at Tyr19 through a dual mechanism: controlling Swe1 kinase stability and Mih1 phosphatase localization. Our findings uncover a novel regulatory network that orchestrates the exit from prophase I, providing new insights into the interplay between recombination surveillance and meiotic cell cycle control.

### Meiotic checkpoint role of Swe1 and Mih1

Previous studies have shown that *SWE1* deletion suppresses the sporulation block of various meiotic mutants, albeit to varying extents ^35,36^, suggesting that Swe1 functions as a target of the MRC. Consistent with these findings, we show that *SWE1* deletion accelerates meiotic progression in *zip1Δ* mutants. However, the *zip1Δ swe1Δ* double mutant still exhibits delayed meiotic divisions compared to the wild type. This contrasts with the nearly complete restoration of normal kinetics of meiotic divisions when checkpoint signaling is abolished by the absence of upstream Mec1 or Mek1 kinases. In addition, Mec1 and Mek1 activity reporters indicate that checkpoint signaling is initially triggered in *zip1Δ swe1Δ*, but it is prematurely downregulated allowing faster and more efficient meiotic progression compared to *zip1Δ*. These findings suggest that Swe1 is not required for checkpoint activation in response to *zip1Δ* defects, but it is essential for its maintenance. Moreover, the disappearance of checkpoint signaling (e.g., Hop1-T318ph) at later time points in *zip1Δ swe1Δ* suggests that the recombination intermediates sensed by the checkpoint in *zip1Δ*, likely unrepaired DSBs ^4,7^, are eventually repaired when high M-CDK activity is restored by the loss of Cdc28-Y19 inhibitory phosphorylation. This conclusion is further supported by the persistence of high levels of Hop1-T318ph and H3-T11ph in *zip1Δ mih1Δ* cells that accumulate Cdc28-Y19ph and exhibit a tighter meiotic block. We propose that, at least in the case of *zip1Δ*, alleviation of the meiotic block by *SWE1* deletion is linked to DSB repair promoted by M-CDK. The moderate spore viability of *zip1Δ swe1Δ* (aprox. 40%) ^35,37^ is consistent with this notion.

Using G1-synchronized cultures, a slight delay in mitotic entry has been observed in vegetative *mih1Δ* cells ^51^. Due to the limited synchrony of our meiotic cultures, we cannot rule out subtle delays, but overall, meiotic progression in the *mih1Δ* mutant appears largely normal. This suggests that the low and transient phosphorylation of Cdc28-Y19 observed in wild-type meiosis can be reversed without Mih1 phosphatase activity, possibly through Swe1 downregulation and/or by the action of an alternative phosphatase ^52,53^ to ensure timely entry into meiosis I. However, Mih1 becomes more critical under conditions where high levels of Cdc28-Y19 phosphorylation arise due to meiotic defects, as seen in *zip1Δ*. Although *zip1Δ* exhibits a strong checkpoint-dependent prophase I arrest, a fraction of cells eventually bypass the arrest and undergo meiotic divisions. The mechanism underlying this checkpoint escape remains unclear, but Mih1 appears to play a role, as the *zip1Δ mih1Δ* double mutant displays more pronounced arrest and sustained inhibitory CDK phosphorylation.

In mitotically dividing cells, Wee1-related kinases are required for cell cycle arrest in response to DNA Damage in various organisms, but in budding yeast Swe1 seems to be dispensable ^33,38,54^. However, more recent reports indicate that Swe1 is indeed involved in the response to genotoxic insults during DNA replication in *S. cerevisiae* ^39,55,56^. It has been reported that, upon hydroxyurea treatment, Mec1 phosphorylates Swe1 at S385 functioning in a parallel pathway to Rad53 (the “mitotic” FHA effector kinase orthologous to the “meiotic” Mek1) to restrain M-CDK activity ^39^. Our results indicate that the situation in the response to meiotic DSBs is different as the *swe1-S385A* mutant does not show a checkpoint defect, and Swe1 acts downstream of Mek1, not in parallel, in the MRC pathway.

### A Novel meiotic role for Cdc5: regulation of Swe1 and Mih1

The observation that Swe1 levels, and consequently Cdc28-Y19 phosphorylation, remain high in *zip1Δ ndt80Δ* cells even when the checkpoint is abrogated by *MEK1* deletion suggests that an Ndt80-dependent factor is required for Swe1 downregulation. The polo-like kinase Cdc5 emerged as a strong candidate for this role, as it functions in multiple events associated with prophase I exit ^22-25,37,57^, coinciding with a decrease in Swe1-mediated inhibitory phosphorylation of CDK ^40^. Supporting this idea, Cdc5 is known to interact with Swe1, and to promote its degradation at the bud neck during the G2/M transition in vegetative cells ^41-44,58^. In line with this, we show that in meiotic prophase I-arrested *zip1Δ ndt80Δ* cells, induction of *CDC5* alone is sufficient to trigger Swe1 degradation and reduce Cdc28-Y19 phosphorylation. Cdc5 has also been reported to promote the degradation of the meiotic chromosome axis protein Red1. Although the precise target(s) of Cdc5 in the SC remains to be determined; this degradation leads to downregulation of Mek1 checkpoint activity ^20^. Consistent with this, we have observed diminished checkpoint signaling upon Cdc5 induction in *zip1Δ ndt80Δ CDC5-IN* cells, as indicated by decreased levels of Hop1-T318ph and H3-T11ph ^59^. These findings raised the possibility that Cdc5-mediated Swe1 reduction may be an indirect consequence of checkpoint inactivation. However, in *zip1Δ ndt80Δ mek1Δ* cells, where checkpoint signaling is absent, but *CDC5* is not expressed due to the absence of Ndt80, Swe1 protein and Cdc28-Y19 phosphorylation persist (Fig. 1c). These findings indicate that Cdc5 promotes Swe1 degradation independently of Mek1 activity, highlighting its direct role in regulating Swe1 stability during meiotic progression.

Both the morphogenesis checkpoint and the MRC rely on Swe1 accumulation to delay entry into mitosis and meiosis I, respectively. In vegetative cells, Swe1 is recruited to the bud neck by the Hsl1 kinase and the Hsl7 adaptor, where it is primed for degradation through the coordinated action of Cla4, Cdc28 and Cdc5 kinases once the morphogenesis checkpoint has been satisfied ^41,44,60,61^. However, in meiosis, Swe1 degradation in meiosis appears to follow a distinct mechanism, despite also involving Cdc5. As meiotic cells lack a bud neck structure, we observed that Swe1 is localized in the nucleus during prophase I, but disappears upon entry into meiosis I coinciding with the normal timing of Cdc5 expression. Similarly, Swe1 nuclear signal is lost in prophase I-arrested *zip1Δ ndt80Δ CDC5-IN* cells upon Cdc5 induction. Notably, we found that, in contrast to the mitotic paradigm, Swe1 degradation in meiosis does not require prior CDK-mediated priming. Furthermore, when Cdc5 and Swe1 are artificially colocalized at an ectopic site outside the nucleus, Cdc5 kinase activity is sufficient to reduce Swe1 levels. These findings strongly suggest that Cdc5 can directly promote Swe1 degradation in meiosis without additional spatial or priming requirements. Consistent with our observations in vivo, Cdc5 can phosphorylate Swe1 in vitro ^44^. Nevertheless, we show here that the retarded gel mobility of Swe1 observed upon *CDC5* expression in prophase I arrested cells does depend on Cdk1 and that, like in mitosis ^60^, Cdk1 activity is required to stimulate Swe1 in meiosis.

Very little was previously known about the regulation of Mih1, the CDK-activating phosphatase, in *S. cerevisiae* meiosis. Our time-lapse microscopy data show that Mih1 is cytoplasmic and excluded from the nucleus during prophase I, but relocates to the nucleus as cells exit prophase I, concurrent with the requirement for M-CDK to drive entry into meiosis I. We further show that this nuclear translocation of Mih1 is dependent on Cdc5. In *S. pombe*, activation of the meiotic recombination checkpoint leads to phosphorylation of the Cdc25 phosphatase by Mek1 and results in its cytoplasmic retention, at least in part, via 14-3-3 protein binding ^62,63^. In *S. cerevisiae*, by contrast, we observe no change in Mih1 localization in the absence of Mek1, indicating a distinct mechanism of checkpoint control. In vegetive cells, Mih1 is dephosphorylated by the PP2A phosphatase during the G2/M transition ^51^. In our system, Mih1 exhibits faster-migrating bands on SDS-PAGE following *CDC5* induction in prophase I-arrested cells, also suggesting Mih1 dephosphorylation. Whereas in human somatic cells phosphorylation of CDC25B/C by PLK1 triggers their nuclear relocation ^64,65^, our results suggest that, in budding yeast meiosis, Cdc5 likely does not directly phosphorylate Mih1 but it may instead activate a phosphatase that promotes Mih1 nuclear entry and/or activation. Although multiple Cdc5-dependent phosphorylation sites have been detected on Mih1 during meiosis ^66^, most of them conform to CDK consensus motifs, hinting at a complex regulatory loop involving an interplay among kinases (Cdc5, Cdk1) and phosphatases (perhaps PP2A) in controlling Mih1 localization and activity.

### Model and concluding remarks

We propose that, at the end of prophase I when meiotic DSBs have been repaired, induction of *CDC5* by Ndt80 contributes to M-CDK activation through a dual mechanism: 1) degradation of the nuclear Swe1 kinase, and 2) translocation of the Mih1 phosphatase into the nucleus (Fig. 6). Together, these processes lead to the elimination of inhibitory phosphorylation on Cdk1 and enable progression into and through meiosis I. In addition, the expression of the major meiosis I B-type cyclin, Clb1, also under Ndt80 control, further amplifies M-Cdk1 activity.

**Fig 6.**
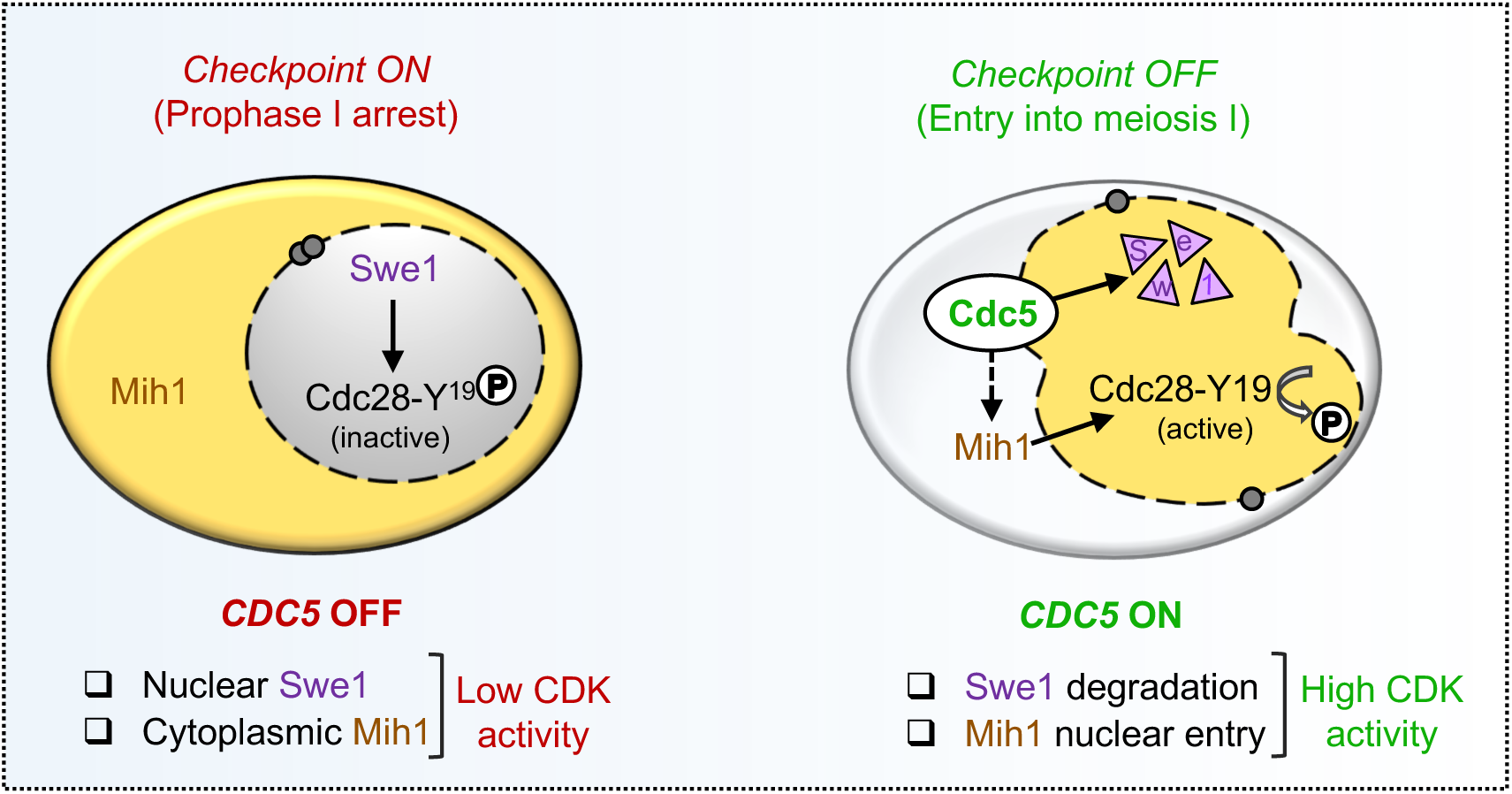
Model for Cdc5-dependent regulation of CDK during the prophase I to meiosis I transition. Dual action of polo-like kinase Cdc5 on Cdc28 (CDK) regulators during budding yeast meiosis. As meiotic recombination is completed and the checkpoint is satisfied, Cdc5 drives degradation of the nuclear Swe1 inhibitory kinase and nuclear translocation of the Mih1 activating phosphatase to promote meiosis I entry by increasing CDK activity. Solid arrow lines imply direct action, whereas discontinuous lines reflect the possibility of an indirect effect. See text for details.

The polo-like kinase family fulfills multiple roles in meiosis across species, from yeasts to mammals, extending beyond the prophase I to meiosis I transition ^67-70^. In mouse oocytes, inhibitory phosphorylation of CDK1 controlled by sequestration of WEE1/MYT1 kinases and CDC25 phosphatases in different subcellular compartments is essential for maintaining the prolonged prophase I arrest. Relocation of these components is required for meiotic reentry ^71,72^. PLK1, the mammalian homolog of Cdc5, is also required for the resumption of meiosis I in oocytes, contributing to the autoamplification loop that promotes full CDK1 activation ^73^. However, the different effects of PLK1 and CDK1 inhibition on oocyte maturation suggest that PLK1 plays additional roles beyond simply activating CDK1 ^74^.

In budding yeast, the essential functions of Cdc5 in promoting exit from prophase I have previously been attributed mainly to its roles in Holliday junction resolution and SC disassembly. Here, we uncover a novel function for Cdc5 in promoting M-CDK activity during the prophase I to meiosis I transition via a dual regulatory action on Swe1 and Mih1. Further work will be required to dissect the precise mechanisms involved in the regulation of Cdk1 inhibitory phosphorylation during meiosis in the context of the complex feedback loops that are known to fine-tune cell cycle progression in mitosis ^52,75-77^, and likely in meiosis as well ^78^.

## METHODS

### Yeast strains and plasmids

Yeast strains genotypes and plasmids used are listed in Table S1 and Table S2, respectively. All the strains are isogenic to the BR1919 background ^79^. Generation of *mek1Δ::kanMX6*, *mec1Δ::KlURA3*, *sml1Δ::kanMX6* ^80^, *swe1Δ::LEU2, mih1Δ::ADE2* ^35^, *cdc28-as1*, *cdc5-N209A* ^37^, *ZIP1-GFP-IN* ^40^, *zip1Δ::LYS2*, *ndt80Δ::LEU2*, and *CDC5-IN* ^59^ gene deletions, mutations or modifications has been previously described. C-terminal tagging of Swe1 with 3HA was performed according to ^81^. For N-terminal tagging of Swe1 with 3MYC, a *delitto perfetto* approach ^82^ was followed using as a donor a PCR fragment obtained from strain YBA179 ^35^ encompassing the N-terminal coding region of *SWE1* containing 3MYC. This technique was also used for the tagging of *SWE1* and *MIH1* with GFP at the corresponding genomic loci, under control of the *HOP1* promoter, using DNA fragments obtained from plasmids pSS439 and pSS265 as donors, respectively. Mutation of Ser385 of Swe1 to alanine to generate *swe1-S385A* was also performed by *delitto perfetto* ^82^. To place *IME1* under control of the *CUP1* promoter, the pYM-N2 plasmid was used ^83^.

To generate plasmid pSS439, a 2.8-kb DNA fragment containing *SWE1* ORF and 3’ UTR region was amplified from genomic DNA and cloned into the *Nhe*I-*Sac*I sites of pSS393 containing *P_HOP1_-GFP-PCH2* ^84^ to remove *PCH2* and placing *GFP-SWE1* under control of the *HOP1* promoter in a centromeric plasmid. To generate pSS441, a 2.1-kb a DNA fragment containing *CDC5* ORF was amplified from plasmid pEM110 [pSS293] ^46^ and cloned into the *Not*I-*Xba*I sites of pSS248 ^40^ to express *GFP-CDC5* under control of the *HOP1* promoter in a multicopy plasmid. Plasmid pSS484 derives from pSS441 introducing *cdc5-N209A* by site-directed mutagenesis. Plasmid pSS449 was constructed by cloning a 4.0-kb *Xho*I-*Sac*I DNA fragment from pSJ906 [pSS398] ^85^ containing *SPC42-mCherry-HIS3MX6* into the same sites of the pRS316 centromeric vector ^86^.

### Meiotic time courses, sporulation efficiency and spore viability

For regular meiotic time courses, BR strains were grown in 3.5 ml of 2xSC medium for 20 - 24 hours (2% Glucose, 0.7% Yeast Nitrogen Base without amino acids, 0.05% Adenine and Complete Supplement Mixture from Formedium at twice the particular concentration indicated by the manufacturer), then transferred to YPDA (2.5 ml) and incubated to saturation for additional 8 hours. Cells were harvested, washed with 2% potassium acetate (KAc), resuspended into 2% KAc (10 ml) and incubated at 30°C with vigorous shaking to induce meiosis and sporulation. Both YPDA and 2% KAc were supplemented with 20 mM adenine and 10 mM uracil. The culture volumes were scaled-up when needed. To increase synchrony in meiotic cultures of BR strains, we modified the procedure described for SK1 in ^49^ based on the control of the expression of the master meiotic regulator *IME1* by the *CUP1* promoter (Figure S5A). BR strains containing *P_CUP1_-IME1* were grown in 2xSC medium for 24 hours, transferred to YPDA with 1% glucose and incubated for 8 hours. Cells were washed and resuspended into 2% KAc. After 12 h of incubation, CuSO_4_ was added at a final concentration of 50 μM. Microscopic observation of cells expressing *ZIP1-GFP* in *P_CUP1_-IME1* cultures reveals that 6 hours after Cu^2+^ addition about 75% of cells are in meiotic prophase ^87^. To score meiotic nuclear divisions, samples were taken at different time points, fixed in 70% Ethanol, washed in PBS and stained with 1 mg/ml DAPI for 15 min. At least 300 cells were counted at each time point. To induce *ZIP1-GFP*, *CDC5* or *cdc5-N209A* expression from the *P_GAL1_* promoter in strains also harboring *GAL4.ER* (*ZIP1-GFP-IN*, *CDC5-IN* or *cdc5-N209A-IN* strains, respectively), 1 μM β-estradiol (Sigma E2257; dissolved in ethanol) was added to the cultures at 24 h in the experiments presented in Fig. 2a-2f, 3b, 4b). To inhibit CDK activity in *CDC5-IN cdc28-as1* strains (Figure 2G), the 1NM-PP1 ATP analog (1 μM), dissolved in DMSO, and 5 μM β-estradiol were added at 24 h. Sporulation efficiency was quantitated by microscopic examination of asci formation after 3 days on sporulation plates. Both mature and immature asci were scored. At least 300 cells were counted for every strain. Spore viability was assessed by tetrad dissection.

### Western blotting

Total cell extracts for Western blot analysis were prepared by trichloroacetic acid (TCA) precipitation from 3-ml aliquots of sporulation cultures. Briefly, cells were washed with cold water and 20% TCA, resuspended in 100 μl of 20% TCA and stored at -80°C. Samples were thawed on ice, and glass beads (0.5 mm diameter) were added to the 0.5 ml mark. Cells were broken using a FastPrep bead beater (MP Biomedicals) with 3 pulses of 15 sec at the 5.0 force setting, keeping the tubes on ice between pulses. Lysates were recovered, glass beads washed with 200 μl 5% TCA, and the pooled extracts were centrifuged at 3000 rpm for 5 min. Pellets were resuspended into 2x Laemmi sample buffer and neutralized by addition of half volume of 2M Tris. Samples were boiled for 5 min and centrifuged at 13000 rpm prior to be loaded into SDS-polyacrylamide gels. The antibodies used are listed in Table S3. Antibodies were diluted into TBS 0.1% Tween containing 5% milk (for Swe1, Cdc5, Pgk1, MYC, HA, GFP and Pgk1), 5% BSA (Bovine Serum Albumin) (for Cdc28-Y19ph), or 0.1% BSA (for Hop1-T318ph). PVDF membranes were incubated at 4°C overnight (primary antibodies), or at room temperature for 1 h (secondary antibodies). The Pierce ECL Plus reagent (ThermoFisher Scientific) was used for detection. The signal was captured with film exposure (Figure 1) or with a Fusion FX6 system (Vilber) (Figures 2-5).

### Microscopy

Images of cells expressing *GFP-SWE1, GFP-CDC5* and *PIL1-GBP-mCherry* shown in Fig. 4a, 4b and S5b were captured with an Olympus IX71 fluorescence microscope equipped with a personal DeltaVision system, a CoolSnap HQ2 (Photometrics) camera, and 100x UPLSAPO 1.4 NA objective. Z-stacks of 13 planes at 0.3-μm intervals were collected in Fig. 4a, b, whereas a single plane was captured in Fig. S5b. Exposure times for GFP-Swe1 were 100 ms (Fig. 4a), or 1000 ms (Fig. S5b). Exposure times for GFP-Cdc5 were 500 ms (Fig. 4b). Exposure times for DIC were 100 ms (Fig. 4a), 50 ms (Fig. 4b) or 80 ms (Fig. S5b), with neutral density filter transmitting 32% of intensity.

For time-lapse experiments, an Andor Dragonfly 200 confocal microscope, equipped with a 100x/1.45 NA Plan Apo objective, and a sCMOS Sona 4.2B-11 camera, controlled by the Fusion 2.2 software was used. 300 μl of prophase I cells were placed on microscopy culture chambers (µ-slide 8 well, Ibidi) previously treated with 0.5 mg/ml of Concanavalin A Type IV (Sigma-Aldrich) and let sit for 5 min. Unfixed cells were removed by pipetting and new 2% KAc medium was added. When needed, β-estradiol or CuSO_4_ were added in the chamber to induce *CDC5* or *IME1* expression, respectively. Chambers were maintained at 30°C throughout the experiments. For the experiments shown in Fig. 3a, and 5a, Z-stacks of 13 planes at 0.4-μm steps were collected at 1 h intervals. Exposure times were 200 ms (488 nm 20% laser power) for GFP-Swe1, 600 ms (488 nm 40% laser power) for GFP-Mih1, 400 ms (561 nm 30% laser power) for Spc42-mCherry, and 100 ms for brightfield. For the experiments shown in Figures 3B and 5B, Z-stacks of 13 planes at 0.4-μm steps were collected at 0, 3 and 6 h after β-estradiol addition to the chamber. Exposure times were 500 ms (488 nm 50% laser power) for GFP-Swe1, 600 ms (488 nm 40% laser power) for GFP-Mih1, 800 ms (561 nm 74% laser power) for Spc42-mCherry, and 100 ms for brightfield. Captured images were processed and analyzed with FIJI software. Brightness and contrast adjustments were equally applied to all entire images in each experiment. Maximum intensity projections of the planes containing signal are presented in Fig. 3a, 3b, 5a, 5b, and single planes in Fig. 4a, 4b, S5b.

### Digital PCR for gene expression analysis

Cells were collected from 1 mL of meiotic cultures, and total RNA was extracted using the RNeasy Mini Kit (Qiagen 74104), following the manufacturer’s instructions. RNA concentration and purity were assessed using a NanoDrop spectrophotometer (Thermo Fisher Scientific). Complementary DNA (cDNA) was synthesized from 2μg of total RNA using the High-Capacity RNA-to-cDNA kit (Applied Biosystems 4387406) and quantified with Qubit (Thermo Fisher Scientific). Digital PCR (dPCR) was performed on an Applied Biosystems QuantStudio Absolute Q platform using TaqMan assays specific for the gene of interest *SWE1* (FAM-labeled probe), and the reference gene *NUP85* (VIC-labeled probe) whose mRNA levels remain constant during meiosis ^88^. Digital PCR reactions were performed using the Absolute Q universal DNA dPCR Master MIX (Thermo Fisher Scientific). Each reaction mixture contained 2μl of master mix, 0.5μl each of SWE1-FAM and NUP85-VIC probes, 1 μl of diluted cDNA (3.5 ng), and water up to 10 μl of final volume. After loading reactions in each well, 15 μl of isolation buffer were also added. Sample partitioning and reaction program was carried out according to the manufacturer’s protocol. Absolute quantification of *SWE1* expression was obtained by normalization to *NUP85* expression. Data were analyzed using QuantStudio Absolute Q Digital PCR System Software v6, with consistent threshold settings applied across all samples. Quality control parameters of digital PCR, including amplitude plots and partition metrics (total partitions, positive partitions, concentration, confidence intervals, precision, and lambda values), are provided in Figure S4.

### Statistics and software

When applicable, to determine the statistical significance of differences, a One-way ANOVA test was used. P-values were calculated with the GraphPad Prism 9.0 and 10.0 software. P<0.05 (*); P<0.01 (**); P<0.001 (***); P<0.0001 (****). GraphPad Prism software was also used for graphical representation. The nature of the error bars in the graphs and the number of biological replicates are indicated in the corresponding figure legend.

## Acknowledgements

We are grateful to Andrés Clemente and Rafa Lucena for fruitful comments and discussions. We also thank Sue Jaspersen, Doug Kellogg, Rafa Lucena, Jesús Carballo and Michael Lichten for reagents. We are indebted to Carmen Castro for her dedication and advice on microscopy time-lapse experiments. We also acknowledge André Espinal and Nahia Hernández-Quitián for initial help in strains construction, and Joao Matos and Rahel Wettstein for sharing unpublished results. Research was supported by grants PID2021-125830NB-I00 (to PSS) and PID2022-136681NB-I00 (to EQ) from Ministry of Science and Innovation (MCIN/AEI/10.13039/501100011033/) and “FEDER Una manera de hacer Europa”, by grant CSI010P23 (to PSS) from the “Junta de Castilla y León” (co-funded by FEDER), and by grant CIDEGENT2020/41 (to EQ) from Generalitat Valenciana.

## Author contributions

SGA: conceptualization, investigation, formal analysis, visualization.

IAG: investigation, formal analysis.

IGT: investigation.

EQ: conceptualization, resources, funding acquisition

BS: conceptualization, supervision, project administration.

PSS: conceptualization, investigation, supervision, funding acquisition, visualization, writing original draft, review and editing.

All authors revised, commented, and approved the manuscript.

## Data availability

All relevant data are within the manuscript and the Supplementary information files. Any other relevant information is available from the corresponding author upon reasonable request.

## SUPPLEMENTARY INFORMATION

### Supplementary figure legends

**Video S1. Nuclear removal of Swe1.**

Time-lapse movie corresponding to the cell shown in Fig. 3a displayed at 1 frame per second.

**Video S2. Nuclear translocation of Mih1.**

Time-lapse movie corresponding to the cell shown in Fig.5a displayed at 1 frame per second.

**Table S1.** Saccharomyces cerevisiae strains.

| Strain | Genotype* | Source |
| --- | --- | --- |
| DP421 | *MATa/MATα leu2-3,112 his4-260 thr1-4 trp1-289 ura3-1 ade2-1 lys2ΔNheI* | PSS Lab |
| DP422 | *zip1Δ::LYS2* | PSS Lab |
| DP714 | *zip1Δ::LYS2 mek1Δ::kanMX6* | PSS Lab |
| DP1157 | *zip1Δ::LYS2 swe1Δ::LEU2* | PSS Lab |
| DP1185 | *P<sub>GAL1</sub>-ZIP1-GFP::TRP1 P<sub>GPD1</sub>-GAL4(848)-ER::URA3/ura3-1* | PSS Lab |
| DP1339 | *zip1Δ::LYS2 swe1-S385A* | This work |
| DP1340 | *SWE1-3HA::kanMX6* | This work |
| DP1341 | *SWE1-3HA::kanMX6 zip1Δ::LYS2* | This work |
| DP1353 | *3MYC-SWE1* | This work |
| DP1353 | *3MYC-SWE1* | This work |
| DP1354 | *3MYC-SWE1 zip1Δ::LYS2* | This work |
| DP1393 | *3MYC-SWE1 zip1Δ::LYS2 mek1Δ::kanMX6* | This work |
| DP1396 | *3MYC-SWE1 zip1Δ::LYS2 ndt80Δ::LEU2 mek1Δ::kanMX6* | This work |
| DP1397 | *3MYC-SWE1 zip1Δ::LYS2 ndt80Δ::LEU2* | This work |
| DP1398 | *3MYC-SWE1 ndt80Δ::LEU2* | This work |
| DP1402 | *mih1Δ::ADE2* | PSS Lab |
| DP1403 | *zip1Δ::LYS2 mih1Δ::ADE2* | PSS Lab |
| DP1404 | *zip1Δ::LYS2 mih1Δ::ADE2 mek1Δ::kanMX6* | This work |
| DP1406 | *zip1Δ::LYS2 swe1-S385A mek1Δ::kanMX6* | This work |
| DP1407 | *3MYC-SWE1 zip1Δ::LYS2 mec1Δ::URA sml1Δ::kanMX6* | This work |
| DP1711 | *zip1Δ::LYS2 ndt80Δ::LEU2 CDC5-IN[=P<sub>CDC5</sub>-CDC5--GAL4(ER)--natMX4--P<sub>GAL1</sub>-CDC5] 3MYC-SWE1* | This work |
| DP1713 | *P<sub>CUP1</sub>-IME1::natNT2* | This work |
| DP1899 | *P<sub>CUP1</sub>-IME1::natNT2 zip1Δ::LYS2 P<sub>HOP1</sub>-GFP-SWE1/swe1Δ::hphMX4* | This work |
| DP1922 | *P<sub>CUP1</sub>-IME1::natNT2 P<sub>HOP1</sub>-GFP-SWE1/SWE1* | This work |
| DP1923 | *P<sub>CUP1</sub>-IME1::natNT2 P<sub>SWE1</sub>-GFP-SWE1/SWE1 zip1Δ::LYS2* | This work |
| DP1924 | *P<sub>CUP1</sub>-IME1::natNT2 P<sub>SWE1</sub>-GFP-SWE1* | This work |
| DP1928 | *P<sub>CUP1</sub>-IME1::natNT2 P<sub>HOP1</sub>-GFP-MIH1/MIH1* | This work |
| DP1929 | *P<sub>CUP1</sub>-IME1::natNT2 P<sub>HOP1</sub>-GFP-MIH1/MIH1 zip1Δ::LYS2* | This work |
| DP1931 | *zip1Δ::LYS2 ndt80Δ::LEU2 CDC5-IN[=P<sub>CDC5</sub>-CDC5--GAL4(ER)--natMX4--P<sub>GAL1</sub>-CDC5] 3MYC-SWE1 P<sub>HOP1</sub>-GFP-MIH1/MIH1* | This work |
| DP1932 | *zip1Δ::LYS2 ndt80Δ::LEU2 CDC5-IN[=P<sub>CDC5</sub>-CDC5--GAL4(ER)--natMX4--P<sub>GAL1</sub>-CDC5] 3MYC-SWE1 cdc28-as1* | This work |
| DP1934 | *zip1Δ::LYS2 ndt80Δ::LEU2 CDC5-IN[=P<sub>CDC5</sub>-CDC5--GAL4(ER)--natMX4--P<sub>GAL1</sub>-CDC5] P<sub>HOP1</sub>-GFP-SWE1/3MYC-SWE1* | This work |
| DP1935 | *P<sub>CUP1</sub>-IME1::natNT2 zip1Δ::LYS2* | This work |
| DP1936 | *zip1Δ::LYS2 ndt80Δ::LEU2 CDC5-IN[=P<sub>CDC5</sub>-CDC5--GAL4(ER)--natMX4--P<sub>GAL1</sub>-CDC5] 3MYC-SWE1 P<sub>HOP1</sub>-GFP-MIH1/MIH1 mek1Δ::kanMX6* | This work |
| DP1977 | *zip1Δ::LYS2 ndt80Δ::LEU2 CDC5-IN[=P<sub>CDC5</sub>-CDC5--GAL4(ER)--natMX4--P<sub>GAL1</sub>-CDC5] 3MYC-SWE1 cdc28-as1 P<sub>HOP1</sub>-GFP-MIH1/MIH1* | This work |
| DP1978 | *P<sub>CUP1</sub>-IME1::natNT2 zip1Δ::LYS2 P<sub>HOP1</sub>-GFP-SWE1/swe1Δ::kanMX6 PIL1-GBP-mCherry::hphMX6/PIL1* | This work |
| DP2121 | *zip1Δ::LYS2 ndt80Δ::LEU2 cdc5-N209A-IN[=P<sub>CDC5</sub>-CDC5--GAL4(ER)--natMX4--P<sub>GAL1</sub>-cdc5-N209A] 3MYC-SWE1* | This work |
| DP2127 | *P<sub>CUP1</sub>-IME1::natNT2 zip1Δ::LYS2 ndt80Δ::LEU2 PIL1-GBP-mCherry::hphMX6/PIL1 P<sub>HOP1</sub>-GFP-SWE1/swe1Δ::hphMX4* | This work |
| DP2133 | *P<sub>CUP1</sub>-IME1::natNT2 zip1Δ::LYS2 PIL1-GBP-mCherry::hphMX6/PIL1* | This work |
| DP2928 | *P<sub>CUP1</sub>-IME1::natNT2 P<sub>HOP1</sub>-GFP-MIH1/MIH1* | This work |

\*All strains are diploids isogenic to BR1919 and, unless specified, homozygous for the indicated markers. DP421 is a *lys2* version of the original BR1919-2N (Rockmill and Roeder, 1990).

**Table S2.** Plasmids.

| Plasmid name | Relevant description | Source/reference |
| --- | --- | --- |
| pRS314 | *CEN6 TRP1* | Sikorski and Hieter, 1989 |
| pRS316 | *CEN6 URA3* | Sikorski and Hieter, 1989 |
| pSS248 | *2μ URA3 P<sub>HOP1</sub>-GFP* | PSS lab<br>González-Arranz et al. (2018) |
| pSS265 | *2μ URA3 P<sub>HOP1</sub>-GFP-MIH1* | PSS lab<br>González-Arranz et al. (2018) |
| pSS439 | *CEN6 TRP1 P<sub>HOP1</sub>-GFP-SWE1* | This work |
| pSS441 | *2μ URA3 P<sub>HOP1</sub>-GFP-CDC5* | This work |
| pSS449 | *CEN6 URA3 SPC42-mCherry-HIS3MX6* | This work |
| pSS484 | *2μ URA3 P<sub>HOP1</sub>-GFP-cdc5-N209A* | This work |

**Table S3.** Antibodies.

| **Antibody** | **Host and type** | **Dilution** | **Source / Reference** |
| --- | --- | --- | --- |
| Cdc2-Y15ph<br>(Cdc28-Y19ph) | Rabbit polyclonal | 1:1000 | Cell Signaling Technology<br>#9111 |
| Cdc5 | Goat polyclonal | 1:1000 | Santa Cruz Biotechnology<br>sc-6733 |
| H3-T11ph | Rabbit polyclonal | 1:5000 | Abcam<br>ab5168 |
| Hop1-T318ph | Rabbit polyclonal | 1:1000 | J. Carballo |
| Ndt80 | Rabbit polyclonal | 1:5000 | M. Lichten |
| Pgk1 (22C5D8) | Mouse monoclonal | 1:5000 | Invitrogene<br>459240 |
| Swe1 | Rabbit polyclonal | 1:5000 | R. Lucena/ D. Kellogg |
| GFP (JL-8) | Mouse monoclonal | 1:1000 | Clontech<br>632381 |
| Myc | Rabbit polyclonal | 1:1000 | Sigma<br>c3956 |
| Anti-goat-HRP | Donkey polyclonal | 1:5000 | Santa Cruz Biotechnology<br>Sc-2033 |
| Anti-mouse-HRP | Sheep polyclonal | 1:5000 | GE-Healthcare<br>NA931 |
| Anti-rabbit-HRP | Donkey polyclonal | 1:5000 | GE-Healthcare<br>NA934 |

**Fig S1.**
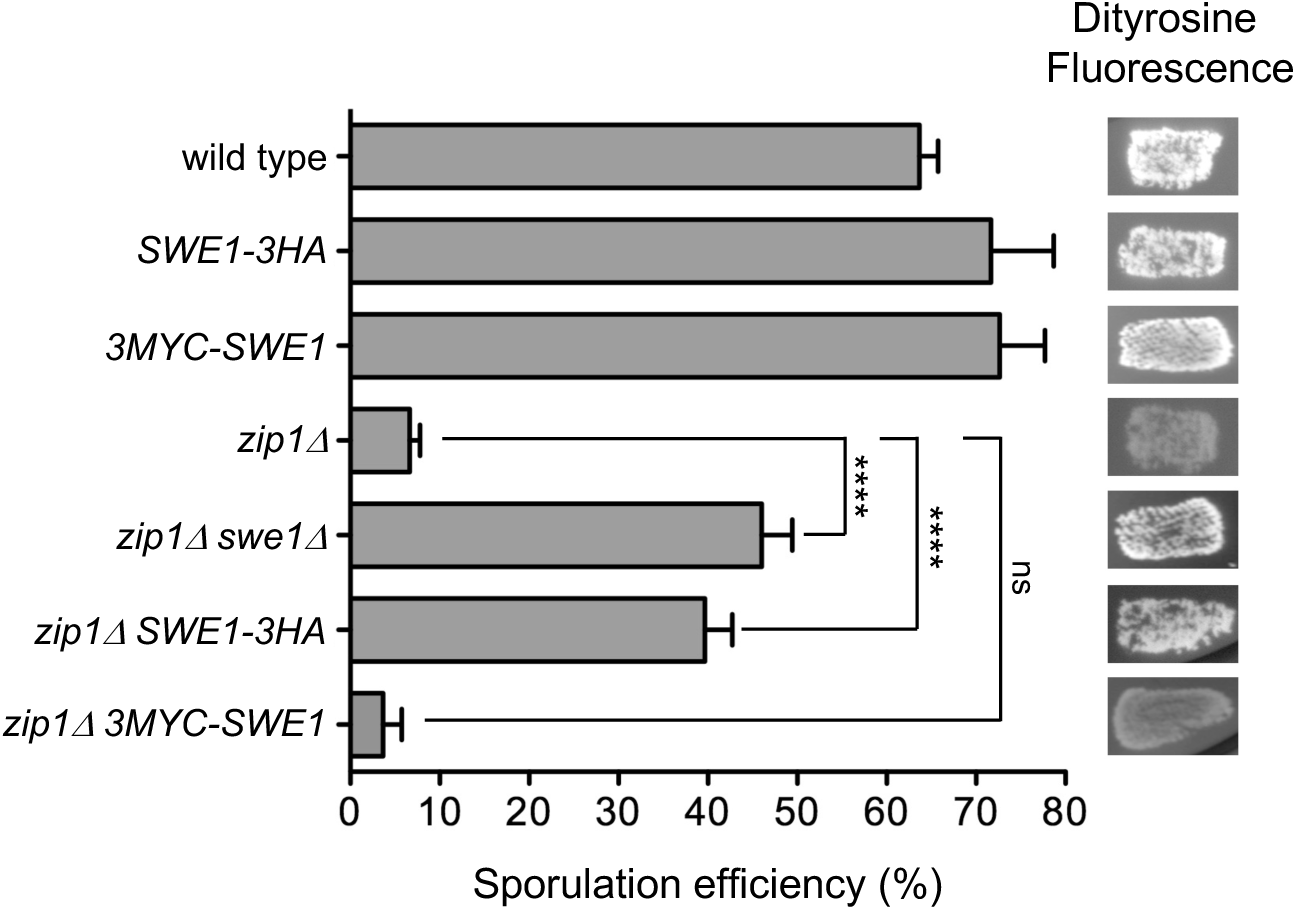
N-terminal tagging of Swe1 with 3MYC supports checkpoint function. Sporulation efficiency and dityrosine fluorescence (DiTyr) were examined after 3 days on sporulation plates. In contrast to *zip1Δ SWE1-3HA*, the *zip1Δ 3MYC-SWE1* strain exhibits low sporulation and reduced DiTyr fluorescence comparable to *zip1Δ*. Error bars, SD; n=3. At least 300 cells were counted for each strain. Strains are DP421 (wild type), DP1340 (*SWE1-3HA*), DP1353 (*3MYC-SWE1*), DP422 (*zip1Δ*), DP1157 (*zip1Δ swe1Δ*), DP1341 (*zip1Δ SWE1-3HA*), and DP1354 (*zip1Δ 3MYC-SWE1*).

**Fig S2.**
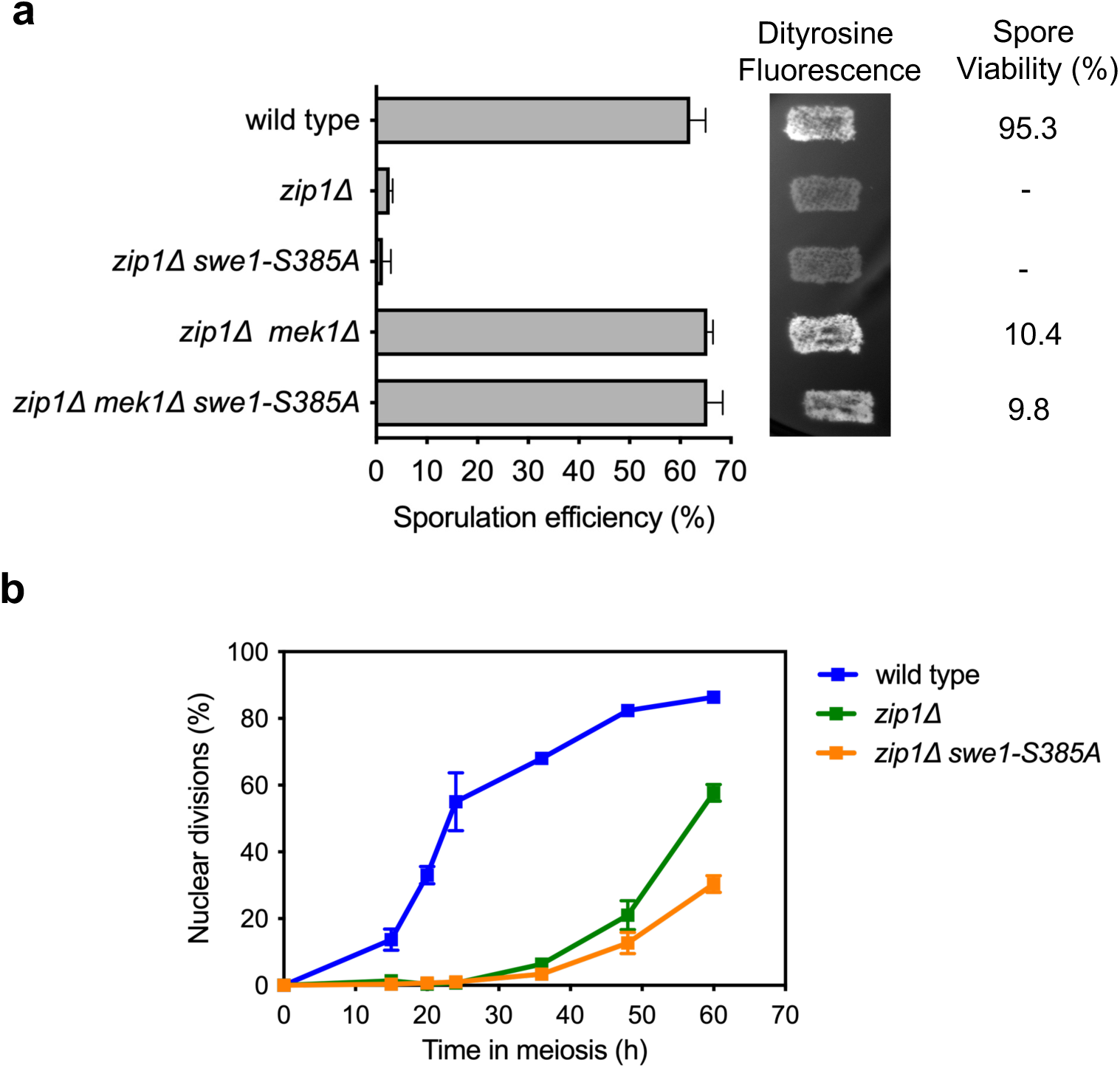
The meiotic recombination checkpoint is functional in the *swe1-S385A* mutant lacking a potential Mec1 phosphorylation site. **(a)** Sporulation efficiency and dityrosine fluorescence (DiTyr) were examined after 3 days on sporulation plates. The *zip1Δ swe1-S385A* double mutant exhibits low sporulation and reduced DiTyr fluorescence comparable to *zip1Δ*. Error bars, SD; n=3. At least 300 cells were counted for each strain. **(b)** Time-course analysis of meiotic nuclear divisions; the percentage of cells containing two or more nuclei is represented. Error bars: SD; n=3. At least 300 cells were scored for each strain at every time point. Strains are DP421 (wild type), DP422 (*zip1Δ*), DP1339 (*zip1Δ swe1-S385A*), DP714 (*zip1Δ mek1Δ*), and DP1406 (*zip1Δ mek1Δ swe1-S385A*).

**Fig S3.**
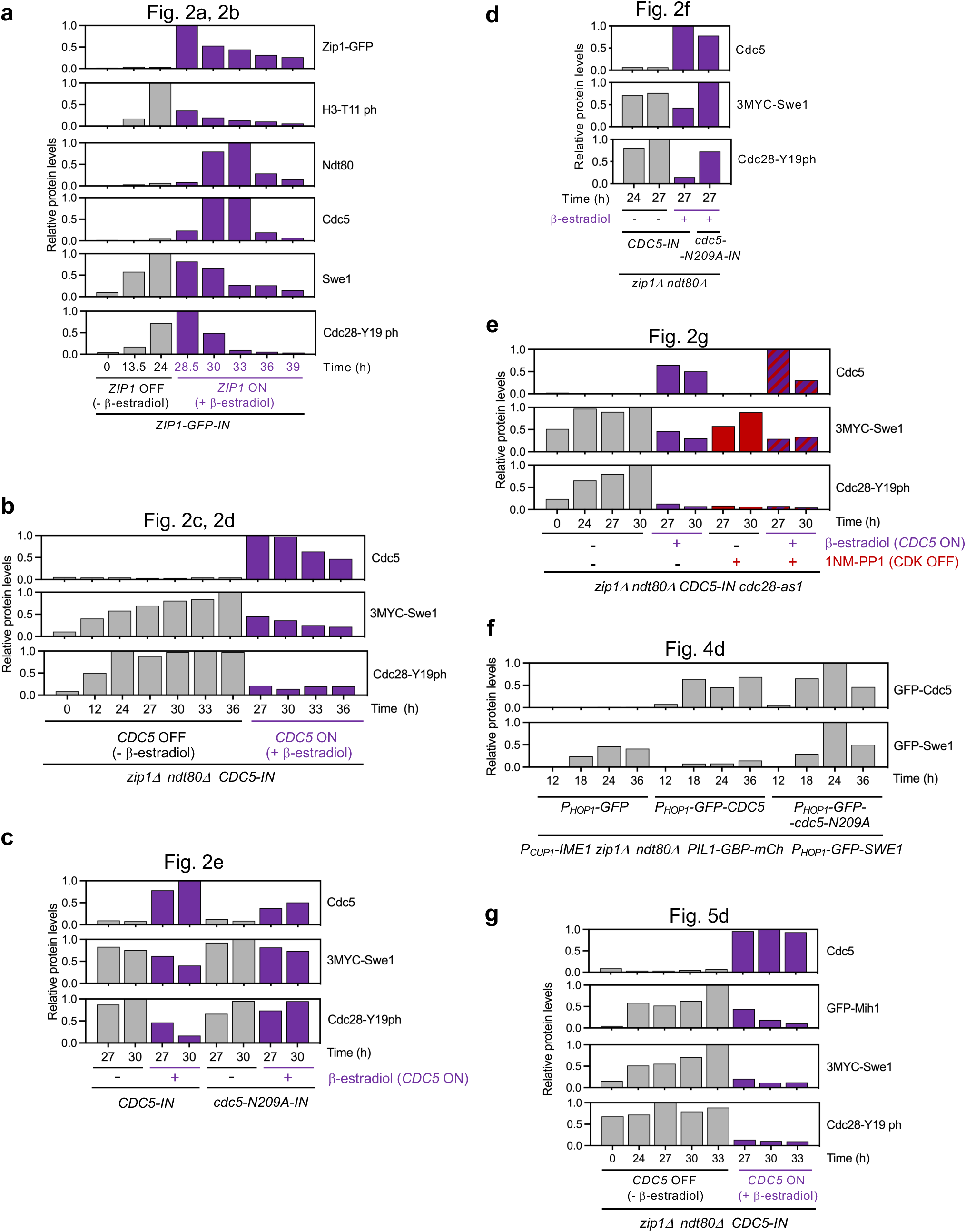
Quantification of relative protein levels in western blot analyses. Protein levels were normalized with Pgk1 and relativized to the maximum value in each experiment, which was set to 1. **(a)** Quantification of Fig 2b. **(b)** Quantification of Fig 2d. **(c)** Quantification of Fig 2e. **(d)** Quantification of Fig 2f. **(e)** Quantification of Fig 2g. **(f)** Quantification of Fig 4d. **(g)** Quantification of Fig 5d.

**Fig. S4.**
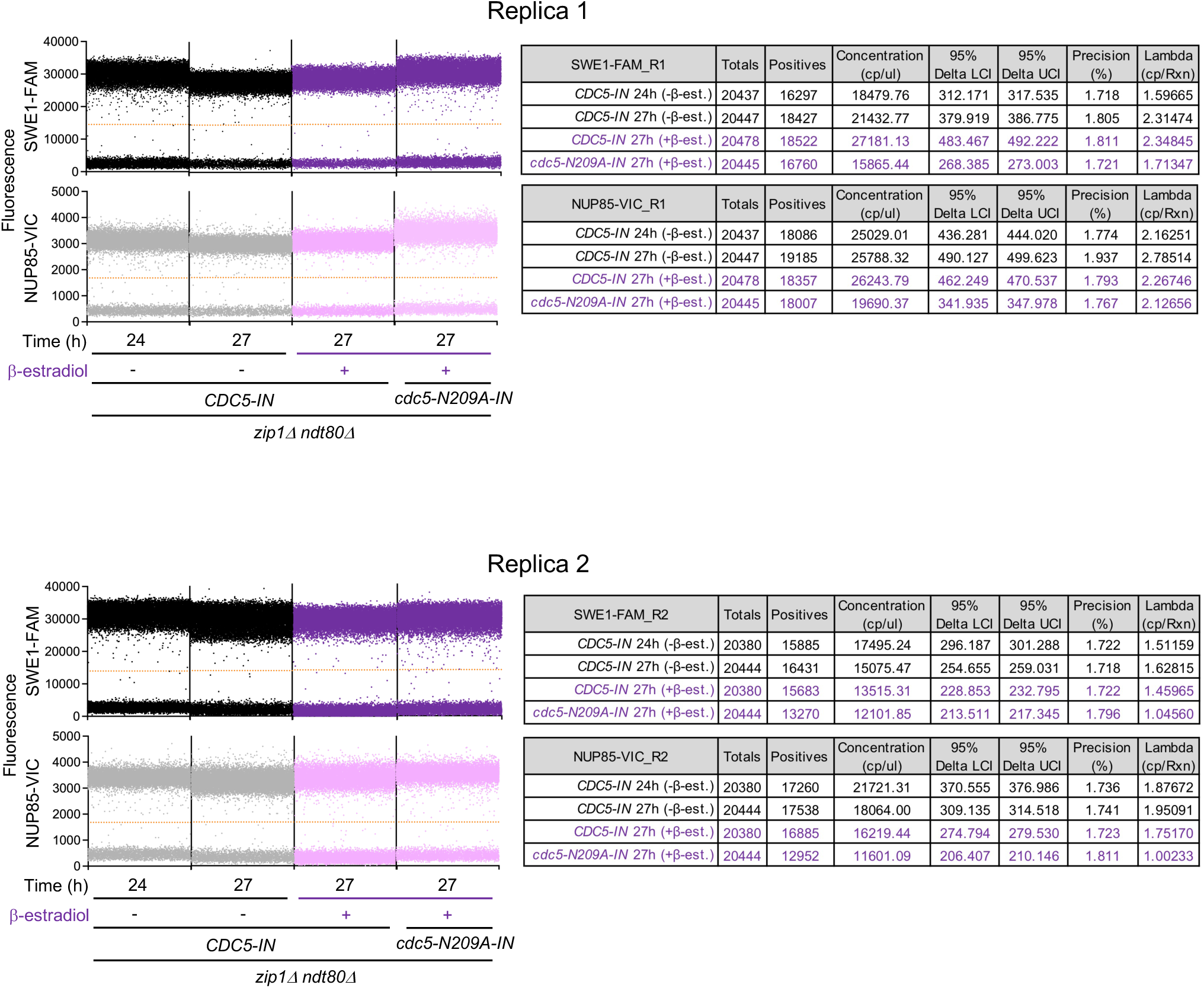
Digital PCR quality control for *SWE1* and *NUP85* expression (related to Fig. 2f). Left panels show amplitude plots for *SWE1* (FAM channel) and the control gene *NUP85* (VIC channel) for all conditions analyzed in the digital PCR gene expression analysis presented in Fig. 2f. Tables on the right summarize key quality metrics obtained, including total partitions, positive partitions, concentration (copies/µL), 95% confidence interval (lower and upper limits), precision (%), and lambda (copies/reaction). Results are presented for both replicates.

**Fig S5.**
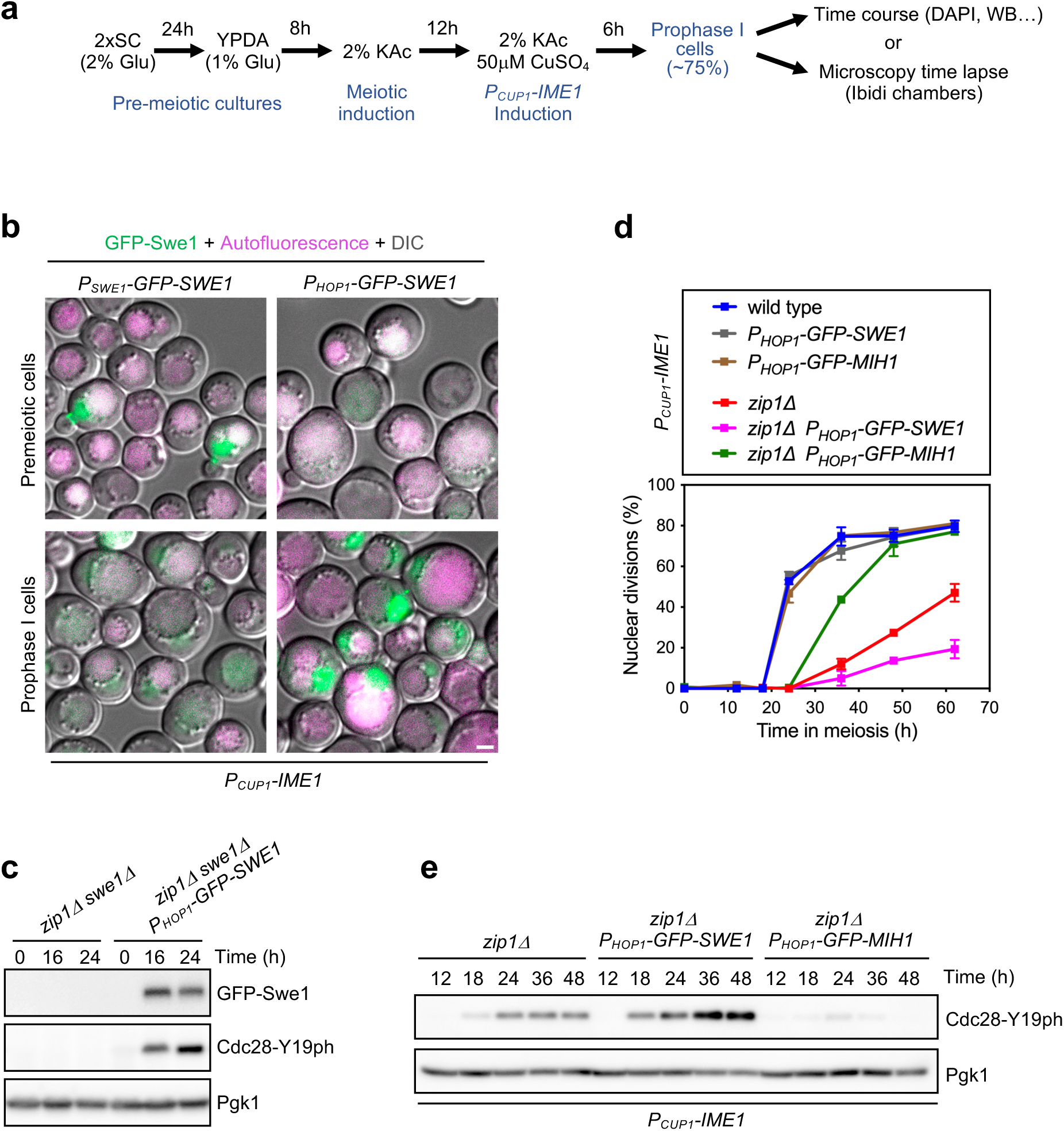
Analysis of GFP-Swe1 and GFP-Mih1 protein function and dynamics. **(a)** Schematic representation of the optimized procedure to increase meiotic synchrony in BR strains based on the induced expression of *IME1* from the *CUP1* promoter. This culture setup was used in the experiments presented in Fig. 3a, 4a, 4b, 4d, 5a, S5b, S5d and S5e. **(b)** Fluorescence microscopy images of GFP-Swe1 in premeiotic cells (top panels), and in meiotic prophase I cells (bottom panels). Expression of *GFP-SWE1* was controlled by the *SWE1* promoter (left panels) or by the *HOP1* promoter (right panels). Synchronized *P_CUP1_-IME1* cultures, as outlined in (a) were used. Samples were taken at 12 h in 2% KAc (premeiotic cells) and 6 h after Cu2^+^ addition; that is, *IME1* induction (prophase I cells). GFP-Swe1 signal is shown in green. Vacuolar background autofluorescence is shown in magenta. Strains are DP1924 (*P_CUP1_-IME1 P_SWE1_-GFP-SWE1*) and DP1922 (*P_CUP1_-IME1 P_HOP1_-GFP-SWE1*). (**c)** Western blot analysis of GFP-Swe1 production and Cdc28-Y19 phosphorylation in the indicated strains. Pgk1 was used as loading control. The strain is DP1157 (*zip1Δ swe1Δ*) transformed with plasmids pRS314 (empty vector) or pSS439 (*P_HOP1_-GFP-SWE1*). **(d)** Time-course analysis of meiotic nuclear divisions; the percentage of cells containing two or more nuclei is represented. Error bars: SD; n=3. At least 300 cells were scored for each strain at every time point. **(e)** Western blot analysis of Cdc28-Y19 phosphorylation. Pgk1 was used as loading control. Strains in (d, e) are DP1713 (*P_CUP1_-IME1*), DP1922 (*P_CUP1_-IME1 P_HOP1_-GFP-SWE1*), DP1928 (*P_CUP1_-IME1 P_HOP1_-GFP-MIH1*), DP1935 (*P_CUP1_-IME1 zip1Δ*), DP1923 (*P_CUP1_-IME1 P_HOP1_-GFP-SWE1 zip1Δ*) and DP1929 (*P_CUP1_-IME1 P_HOP1_-GFP-MIH1 zip1Δ*). DP1922, DP1928, DP1923 and DP1929 also contained pSS449 (*SPC42-mCherry*).

## REFERENCES

1 Keeney, S., Lange, J. & Mohibullah, N. Self-organization of meiotic recombination initiation: general principles and molecular pathways. Annu Rev Genet 48, 187–214, doi:10.1146/annurev-genet-120213-092304 (2014).

2 Hunter, N. Meiotic Recombination: The essence of heredity. Cold Spring Harb Perspect Biol 7, a016618, doi:10.1101/cshperspect.a016618 (2015).

3 Borner, G. V., Hochwagen, A. & MacQueen, A. J. Meiosis in budding yeast. Genetics 225, doi:10.1093/genetics/iyad125 (2023).

4 Raghavan, A. R. & Hochwagen, A. Keeping it safe: control of meiotic chromosome breakage. Trends Genet, doi:10.1016/j.tig.2024.11.006 (2024).

5 Subramanian, V. V. & Hochwagen, A. The meiotic checkpoint network: step-by-step through meiotic prophase. Cold Spring Harb Perspect Biol 6, a016675, doi:10.1101/cshperspect.a016675 (2014).

6 Herruzo, E. et al. The Pch2 AAA+ ATPase promotes phosphorylation of the Hop1 meiotic checkpoint adaptor in response to synaptonemal complex defects. Nucleic Acids Res 44, 7722–7741, doi:10.1093/nar/gkw506 (2016).

7 Hollingsworth, N. M. & Gaglione, R. The meiotic-specific Mek1 kinase in budding yeast regulates interhomolog recombination and coordinates meiotic progression with double-strand break repair. Curr Genet 65, 631–641, doi:10.1007/s00294-019-00937-3 (2019).

8 Tung, K. S. & Roeder, G. S. Meiotic chromosome morphology and behavior in zip1 mutants of Saccharomyces cerevisiae. Genetics 149, 817–832, doi:10.1093/genetics/149.2.817 (1998).

9 Bishop, D. K., Park, D., Xu, L. & Kleckner, N. DMC1: a meiosis-specific yeast homolog of *E. coli* recA required for recombination, synaptonemal complex formation, and cell cycle progression. Cell 69, 439–456, doi:10.1016/0092-8674(92)90446-j (1992).

10 Refolio, E., Cavero, S., Marcon, E., Freire, R. & San-Segundo, P. A. The Ddc2/ATRIP checkpoint protein monitors meiotic recombination intermediates J Cell Sci 124, 2488–2500, doi:10.1242/jcs.081711 (2011).

11 Sampathkumar, A. et al. Replication protein-A, RPA, plays a pivotal role in the maintenance of recombination checkpoint in yeast meiosis. Sci Rep 14, 9550, doi:10.1038/s41598-024-60082-x (2024).

12 Hong, E. J. & Roeder, G. S. A role for Ddc1 in signaling meiotic double-strand breaks at the pachytene checkpoint. Genes Dev 16, 363–376, doi:10.1101/gad.938102 (2002).

13 Lydall, D., Nikolsky, Y., Bishop, D. K. & Weinert, T. A meiotic recombination checkpoint controlled by mitotic checkpoint genes. Nature 383, 840–843, doi:10.1038/383840a0 (1996).

14 Carballo, J. A., Johnson, A. L., Sedgwick, S. G. & Cha, R. S. Phosphorylation of the axial element protein Hop1 by Mec1/Tel1 ensures meiotic interhomolog recombination. Cell 132, 758–770, doi:10.1016/j.cell.2008.01.035 (2008).

15 Chuang, C. N., Cheng, Y. H. & Wang, T. F. Mek1 stabilizes Hop1-Thr318 phosphorylation to promote interhomolog recombination and checkpoint responses during yeast meiosis. Nucleic Acids Res 40, 11416–11427, doi:10.1093/nar/gks920 (2012).

16 Niu, H. et al. Partner choice during meiosis is regulated by Hop1-promoted dimerization of Mek1. Mol Biol Cell 16, 5804–5818, doi:10.1091/mbc.e05-05-0465. (2005).

17 Wan, L., de los Santos, T., Zhang, C., Shokat, K. & Hollingsworth, N. M. Mek1 kinase activity functions downstream of RED1 in the regulation of meiotic double strand break repair in budding yeast. Mol Biol Cell 15, 11–23, doi:10.1091/mbc.e03-07-0499 (2004).

18 Chen, X. et al. Mek1 coordinates meiotic progression with DNA break repair by directly phosphorylating and inhibiting the yeast pachytene exit regulator Ndt80. PLoS Genet 14, e1007832, doi:10.1371/journal.pgen.1007832 (2018).

19 Hepworth, S. R., Friesen, H. & Segall, J. *NDT80* and the meiotic recombination checkpoint regulate expression of middle sporulation-specific genes in *Saccharomyces cerevisiae*. Mol Cell Biol 18, 5750–5761, doi:10.1128/MCB.18.10.5750 (1998).

20 Prugar, E., Burnett, C., Chen, X. & Hollingsworth, N. M. Coordination of double strand break repair and meiotic progression in yeast by a Mek1-Ndt80 negative feedback loop. Genetics 206, 497–512, doi:10.1534/genetics.117.199703 (2017).

21 Tung, K. S., Hong, E. J. & Roeder, G. S. The pachytene checkpoint prevents accumulation and phosphorylation of the meiosis-specific transcription factor Ndt80. Proc Natl Acad Sci U S A 97, 12187–12192, doi:10.1073/pnas.220464597 (2000).

22 Argunhan, B. et al. Fundamental cell cycle kinases collaborate to ensure timely destruction of the synaptonemal complex during meiosis. EMBO J 36, 2488–2509, doi:10.15252/embj.201695895 (2017).

23 Matos, J., Blanco, M. G., Maslen, S., Skehel, J. M. & West, S. C. Regulatory control of the resolution of DNA recombination intermediates during meiosis and mitosis. Cell 147, 158–172, doi:10.1016/j.cell.2011.08.032 (2011).

24 Sanchez, A. et al. Exo1 recruits Cdc5 polo kinase to MutLgamma to ensure efficient meiotic crossover formation. Proc Natl Acad Sci U S A 117, 30577–30588, doi:10.1073/pnas.2013012117 (2020).

25 Sourirajan, A. & Lichten, M. Polo-like kinase Cdc5 drives exit from pachytene during budding yeast meiosis. Genes Dev 22, 2627–2632, doi:10.1101/gad.1711408 (2008).

26 MacKenzie, A. M. & Lacefield, S. CDK Regulation of Meiosis: Lessons from S. cerevisiae and S. pombe. Genes 11, doi:10.3390/genes11070723 (2020).

27 Palacios-Blanco, I. & Martin-Castellanos, C. Cyclins and CDKs in the regulation of meiosis-specific events. Front Cell Dev Biol 10, 1069064, doi:10.3389/fcell.2022.1069064 (2022).

28 Carlile, T. M. & Amon, A. Meiosis I is established through division-specific translational control of a cyclin. Cell 133, 280–291, doi:10.1016/j.cell.2008.02.032 (2008).

29 Grandin, N. & Reed, S. I. Differential function and expression of Saccharomyces cerevisiae B-type cyclins in mitosis and meiosis. Mol Cell Biol 13, 2113–2125, doi:10.1128/mcb.13.4.2113-2125.1993 (1993).

30 Booher, R. N., Deshaies, R. J. & Kirschner, M. W. Properties of *Saccharomyces cerevisiae* wee1 and its differential regulation of p34CDC28 in response to G1 and G2 cyclins. EMBO J 12, 3417–3426, doi:10.1002/j.1460-2075.1993.tb06016.x (1993).

31 Ma, X. J., Lu, Q. & Grunstein, M. A search for proteins that interact genetically with histone H3 and H4 amino termini uncovers novel regulators of the Swe1 kinase in *Saccharomyces cerevisiae*. Genes Dev 10, 1327–1340, doi:10.1101/gad.10.11.1327 (1996).

32 Russell, P., Moreno, S. & Reed, S. I. Conservation of mitotic controls in fission and budding yeasts. Cell 57, 295–303, doi:10.1016/0092-8674(89)90967-7 (1989).

33 Amon, A., Surana, U., Muroff, I. & Nasmyth, K. Regulation of p34CDC28 tyrosine phosphorylation is not required for entry into mitosis in *S. cerevisiae*. Nature 355, 368–371, doi:10.1038/355368a0 (1992).

34 Sia, R. A., Herald, H. A. & Lew, D. J. Cdc28 tyrosine phosphorylation and the morphogenesis checkpoint in budding yeast. Mol Biol Cell 7, 1657–1666, doi:10.1091/mbc.7.11.1657 (1996).

35 Leu, J. Y. & Roeder, G. S. The pachytene checkpoint in *S. cerevisiae* depends on Swe1-mediated phosphorylation of the cyclin-dependent kinase Cdc28. Mol Cell 4, 805–814, doi:10.1016/s1097-2765(00)80390-1 (1999).

36 Pak, J. & Segall, J. Role of Ndt80, Sum1, and Swe1 as targets of the meiotic recombination checkpoint that control exit from pachytene and spore formation in Saccharomyces cerevisiae. Mol Cell Biol 22, 6430–6440, doi:10.1128/MCB.22.18.6430-6440.2002 (2002).

37 Acosta, I., Ontoso, D. & San-Segundo, P. A. The budding yeast polo-like kinase Cdc5 regulates the Ndt80 branch of the meiotic recombination checkpoint pathway. Mol Biol Cell 22, 3478–3490 (2011).

38 Sorger, P. K. & Murray, A. W. S-phase feedback control in budding yeast independent of tyrosine phosphorylation of p34cdc28. Nature 355, 365–368, doi:10.1038/355365a0 (1992).

39 Palou, G. et al. Three Different Pathways Prevent Chromosome Segregation in the Presence of DNA Damage or Replication Stress in Budding Yeast. PLoS Genet 11, e1005468, doi:10.1371/journal.pgen.1005468 (2015).

40 González-Arranz, S. et al. Functional Impact of the H2A.Z Histone Variant During Meiosis in *Saccharomyces cerevisiae*. Genetics 209, 997–1015, doi:10.1534/genetics.118.301110 genetics.118.301110 [pii] (2018).

41 Asano, S. et al. Concerted mechanism of Swe1/Wee1 regulation by multiple kinases in budding yeast. EMBO J 24, 2194–2204, doi:10.1038/sj.emboj.7600683 (2005).

42 Howell, A. S. & Lew, D. J. Morphogenesis and the cell cycle. Genetics 190, 51–77, doi:10.1534/genetics.111.128314 (2012).

43 Lee, K. S., Asano, S., Park, J. E., Sakchaisri, K. & Erikson, R. L. Monitoring the cell cycle by multi-kinase-dependent regulation of Swe1/Wee1 in budding yeast. Cell Cycle 4, 1346–1349, doi:10.4161/cc.4.10.2049 (2005).

44 Sakchaisri, K. et al. Coupling morphogenesis to mitotic entry. Proc Natl Acad Sci U S A 101, 4124–4129, doi:10.1073/pnas.0400641101 (2004).

45 Elia, A. E., Cantley, L. C. & Yaffe, M. B. Proteomic screen finds pSer/pThr-binding domain localizing Plk1 to mitotic substrates. Science 299, 1228–1231, doi:10.1126/science.1079079 (2003).

46 Mortensen, E. M., Haas, W., Gygi, M., Gygi, S. P. & Kellogg, D. R. Cdc28-dependent regulation of the Cdc5/Polo kinase. Curr Biol 15, 2033–2037, doi:10.1016/j.cub.2005.10.046 (2005).

47 Rodriguez-Rodriguez, J. A., Moyano, Y., Jativa, S. & Queralt, E. Mitotic Exit Function of Polo-like Kinase Cdc5 Is Dependent on Sequential Activation by Cdk1. Cell Rep 15, 2050–2062, doi:10.1016/j.celrep.2016.04.079 (2016).

48 Ubersax, J. A. et al. Targets of the cyclin-dependent kinase Cdk1. Nature 425, 859–864, doi:10.1038/nature02062 (2003).

49 Chia, M. & van Werven, F. J. Temporal Expression of a Master Regulator Drives Synchronous Sporulation in Budding Yeast. G3 6, 3553–3560, doi:10.1534/g3.116.034983 (2016).

50 Herruzo, E. et al. Pch2 orchestrates the meiotic recombination checkpoint from the cytoplasm. PLoS Genet 17, e1009560, doi:10.1371/journal.pgen.1009560 (2021).

51 Pal, G., Paraz, M. T. & Kellogg, D. R. Regulation of Mih1/Cdc25 by protein phosphatase 2A and casein kinase 1. J Cell Biol 180, 931–945, doi:10.1083/jcb.200711014 (2008).

52 Harvey, S. L. et al. A phosphatase threshold sets the level of Cdk1 activity in early mitosis in budding yeast. Mol Biol Cell 22, 3595–3608, doi:10.1091/mbc.E11-04-0340 (2011).

53 Kennedy, E. K. et al. Redundant Regulation of Cdk1 Tyrosine Dephosphorylation in *Saccharomyces cerevisiae*. Genetics 202, 903–910, doi:10.1534/genetics.115.182469 (2016).

54 Stueland, C. S., Lew, D. J., Cismowski, M. J. & Reed, S. I. Full activation of p34CDC28 histone H1 kinase activity is unable to promote entry into mitosis in checkpoint-arrested cells of the yeast *Saccharomyces cerevisiae*. Mol Cell Biol 13, 3744–3755, doi:10.1128/mcb.13.6.3744-3755.1993 (1993).

55 Edenberg, E. R., Mark, K. G. & Toczyski, D. P. Ndd1 turnover by SCF(Grr1) is inhibited by the DNA damage checkpoint in *Saccharomyces cerevisiae*. PLoS Genet 11, e1005162, doi:10.1371/journal.pgen.1005162 (2015).

56 Raspelli, E., Cassani, C., Lucchini, G. & Fraschini, R. Budding yeast Dma1 and Dma2 participate in regulation of Swe1 levels and localization. Mol Biol Cell 22, 2185–2197, doi:10.1091/mbc.E11-02-0127 (2011).

57 Challa, K. et al. Meiosis-specific prophase-like pathway controls cleavage-independent release of cohesin by Wapl phosphorylation. PLoS Genet 15, e1007851, doi:10.1371/journal.pgen.1007851 (2019).

58 Bartholomew, C. R., Woo, S. H., Chung, Y. S., Jones, C. & Hardy, C. F. Cdc5 interacts with the Wee1 kinase in budding yeast. Mol Cell Biol 21, 4949–4959, doi:10.1128/MCB.21.15.4949-4959.2001 (2001).

59 Gonzalez-Arranz, S., Acosta, I., Carballo, J. A., Santos, B. & San-Segundo, P. A. The N-Terminal Region of the Polo Kinase Cdc5 Is Required for Downregulation of the Meiotic Recombination Checkpoint. Cells 10, doi:10.3390/cells10102561 (2021).

60 Harvey, S. L., Charlet, A., Haas, W., Gygi, S. P. & Kellogg, D. R. Cdk1-dependent regulation of the mitotic inhibitor Wee1. Cell 122, 407–420, doi:10.1016/j.cell.2005.05.029 (2005).

61 McMillan, J. N., Theesfeld, C. L., Harrison, J. C., Bardes, E. S. & Lew, D. J. Determinants of Swe1p degradation in *Saccharomyces cerevisiae*. Mol Biol Cell 13, 3560–3575, doi:10.1091/mbc.e02-05-0283 (2002).

62 Perez-Hidalgo, L., Moreno, S. & San-Segundo, P. A. Regulation of meiotic progression by the meiosis-specific checkpoint kinase Mek1 in fission yeast. J Cell Sci 116, 259–271, doi:10.1242/jcs.00232 (2003).

63 Perez-Hidalgo, L., Moreno, S. & San-Segundo, P. A. The fission yeast meiotic checkpoint kinase Mek1 regulates nuclear localization of Cdc25 by phosphorylation. Cell Cycle 7, 3720–3730, doi:10.4161/cc.7.23.7177 (2008).

64 Lobjois, V., Jullien, D., Bouche, J. P. & Ducommun, B. The polo-like kinase 1 regulates CDC25B-dependent mitosis entry. Biochim Biophys Acta 1793, 462–468, doi:10.1016/j.bbamcr.2008.12.015 (2009).

65 Toyoshima-Morimoto, F., Taniguchi, E. & Nishida, E. Plk1 promotes nuclear translocation of human Cdc25C during prophase. EMBO Rep 3, 341–348, doi:10.1093/embo-reports/kvf069 (2002).

66 King, G. A. et al. Meiotic nuclear pore complex remodeling provides key insights into nuclear basket organization. J Cell Biol 222, doi:10.1083/jcb.202204039 (2023).

67 Attner, M. A., Miller, M. P., Ee, L. S., Elkin, S. K. & Amon, A. Polo kinase Cdc5 is a central regulator of meiosis I. Proc Natl Acad Sci U S A 110, 14278–14283, doi:10.1073/pnas.1311845110 (2013).

68 Kalous, J. & Aleshkina, D. Multiple Roles of PLK1 in Mitosis and Meiosis. Cells 12, doi:10.3390/cells12010187 (2023).

69 Bhalla, N. Context is everything: The role of polo-like kinase I during C. elegans oocyte meiosis. J Cell Biol 224, doi:10.1083/jcb.202506208 (2025).

70 Taylor, S. J. P. et al. BUB-1 and CENP-C recruit PLK-1 to control chromosome alignment and segregation during meiosis I in C. elegans oocytes. Elife 12, doi:10.7554/eLife.84057 (2023).

71 Adhikari, D. et al. Inhibitory phosphorylation of Cdk1 mediates prolonged prophase I arrest in female germ cells and is essential for female reproductive lifespan. Cell Res 26, 1212–1225, doi:10.1038/cr.2016.119 (2016).

72 Oh, J. S., Han, S. J. & Conti, M. Wee1B, Myt1, and Cdc25 function in distinct compartments of the mouse oocyte to control meiotic resumption. J Cell Biol 188, 199–207, doi:10.1083/jcb.200907161 (2010).

73 Pahlavan, G. et al. Characterization of polo-like kinase 1 during meiotic maturation of the mouse oocyte. Dev Biol 220, 392–400, doi:10.1006/dbio.2000.9656 (2000).

74 Solc, P. et al. Multiple requirements of PLK1 during mouse oocyte maturation. PLoS One 10, e0116783, doi:10.1371/journal.pone.0116783 (2015).

75 Ferrell, J. E., Jr. Feedback regulation of opposing enzymes generates robust, all-or-none bistable responses. Curr Biol 18, R244–245, doi:10.1016/j.cub.2008.02.035 (2008).

76 Hutter, L. H., Rata, S., Hochegger, H. & Novak, B. Interlinked bistable mechanisms generate robust mitotic transitions. Cell Cycle 16, 1885–1892, doi:10.1080/15384101.2017.1371885 (2017).

77 King, K., Kang, H., Jin, M. & Lew, D. J. Feedback control of Swe1p degradation in the yeast morphogenesis checkpoint. Mol Biol Cell 24, 914–922, doi:10.1091/mbc.E12-11-0812 (2013).

78 Lemonnier, T., Dupre, A. & Jessus, C. The G2-to-M transition from a phosphatase perspective: a new vision of the meiotic division. Cell Div 15, 9, doi:10.1186/s13008-020-00065-2 (2020).

79 Rockmill, B. & Roeder, G. S. Meiosis in asynaptic yeast. Genetics 126, 563–574 (1990).

80 Ontoso, D., Acosta, I., van Leeuwen, F., Freire, R. & San-Segundo, P. A. Dot1-dependent histone H3K79 methylation promotes activation of the Mek1 meiotic checkpoint effector kinase by regulating the Hop1 adaptor. PLoS Genet 9, e1003262, doi:10.1371/journal.pgen.1003262 (2013).

81 Longtine, M. S. et al. Additional modules for versatile and economical PCR-based gene deletion and modification in *Saccharomyces cerevisiae*. Yeast 14, 953–961 (1998).

82 Stuckey, S., Mukherjee, K. & Storici, F. In vivo site-specific mutagenesis and gene collage using the delitto perfetto system in yeast *Saccharomyces cerevisiae*. Methods Mol Biol 745, 173–191, doi:10.1007/978-1-61779-129-1_11 (2011).

83 Janke, C. et al. A versatile toolbox for PCR-based tagging of yeast genes: new fluorescent proteins, more markers and promoter substitution cassettes. Yeast 21, 947–962, doi:10.1002/yea.1142 (2004).

84 Herruzo, E., Santos, B., Freire, R., Carballo, J. A. & San-Segundo, P. A. Characterization of Pch2 localization determinants reveals a nucleolar-independent role in the meiotic recombination checkpoint. Chromosoma 128, 297–316, doi:10.1007/s00412-019-00696-7 (2019).

85 Casey, A. K. et al. Integrity and function of the *Saccharomyces cerevisiae* spindle pole body depends on connections between the membrane proteins Ndc1, Rtn1, and Yop1. Genetics 192, 441–455, doi:10.1534/genetics.112.141465 (2012).

86 Sikorski, R. S. & Hieter, P. A system of shuttle vectors and yeast host strains designed for efficient manipulation of DNA in *Saccharomyces cerevisiae*. Genetics 122, 19–27, doi:10.1093/genetics/122.1.19 (1989).

87 González-Arranz, S. et al. SWR1-independent association of H2A.Z to the LINC complex promotes meiotic chromosome motion. Front Cell Dev Biol 8, 594092, doi:10.3389/fcell.2020.594092 (2020).

88 Govin, J. et al. Systematic screen reveals new functional dynamics of histones H3 and H4 during gametogenesis. Genes Dev 24, 1772–1786, doi:10.1101/gad.1954910 (2010).

## References

Rockmill, B., and G.S. Roeder. 1990. Meiosis in asynaptic yeast. Genetics. 126:563–574.

González-Arranz, S. et al. Functional Impact of the H2A.Z Histone Variant During Meiosis in *Saccharomyces cerevisiae*. Genetics 209, 997–1015 (2018)

Sikorski, R. S. & Hieter, P. A system of shuttle vectors and yeast host strains designed for efficient manipulation of DNA in *Saccharomyces cerevisiae*. Genetics 122, 19–27 (1989).

